# Input zone-selective dysrhythmia in motor thalamus after dopamine depletion

**DOI:** 10.1101/2021.08.30.458172

**Authors:** Kouichi C. Nakamura, Andrew Sharott, Takuma Tanaka, Peter J. Magill

**Author notes:** Correspondence should be addressed to either Dr. Kouichi C. Nakamura or Dr. Peter J. Magill, MRC Brain Network Dynamics Unit, University of Oxford, Mansfield Road, Oxford, OX1 3TH, UK. or.

## Abstract

The cerebral cortex, basal ganglia and motor thalamus form circuits important for purposeful movement. In Parkinsonism, basal ganglia neurons often exhibit dysrhythmic activity during, and with respect to, the slow (∼1 Hz) and beta-band (15–30 Hz) oscillations that emerge in cortex in a brain state-dependent manner. There remains, however, a pressing need to elucidate the extent to which motor thalamus activity becomes similarly dysrhythmic after dopamine depletion relevant to Parkinsonism. To address this, we recorded single-neuron and ensemble outputs in the ‘basal ganglia-recipient zone’ (BZ) and ‘cerebellar-recipient zone’ (CZ) of motor thalamus in anesthetized male dopamine-intact rats and 6-OHDA-lesioned rats during two brain states, respectively defined by cortical slow-wave activity and activation. Two forms of thalamic input zone-selective dysrhythmia manifested after dopamine depletion: First, BZ neurons, but not CZ neurons, exhibited abnormal phase-shifted firing with respect to cortical slow oscillations prevalent during slow-wave activity; secondly, BZ neurons, but not CZ neurons, inappropriately synchronized their firing and engaged with the exaggerated cortical beta oscillations arising in activated states. These dysrhythmias were not accompanied by the thalamic hypoactivity predicted by canonical firing rate-based models of circuit organization in Parkinsonism. Complementary recordings of neurons in substantia nigra pars reticulata suggested their altered activity dynamics could underpin the BZ dysrhythmias. Finally, pharmacological perturbations demonstrated that ongoing activity in the motor thalamus bolsters exaggerated beta oscillations in motor cortex. We conclude that BZ neurons are selectively primed to mediate the detrimental influences of abnormal slow and beta-band rhythms on circuit information processing in Parkinsonism.

## Introduction

Chronic depletion of dopamine from basal ganglia (BG) circuits, as occurs in Parkinson’s disease (PD), profoundly alters the electrical activities of neurons therein. Disturbed BG outputs should have detrimental consequences for their target neurons in the so-called motor thalamus (DeLong, 1990; Rubin et al., 2012; Bosch-Bouju et al., 2013). Because motor thalamic neurons are key effectors of BG outputs, some behavioral impairments in PD likely stem from their aberrant activity dynamics.

Dysrhythmic neuronal activity, *i.e.* abnormal oscillatory firing, is common in basal ganglia-thalamocortical circuits in Parkinsonism (Hammond et al., 2007; Galvan et al., 2015). Neuronal dysrhythmia manifests during sleep and waking states (as well as in general anesthesia), across oscillations with markedly different frequencies. Two exemplary dysrhythmias are associated with slow oscillations (∼1 Hz) and beta-band oscillations (15–30 Hz). After dopamine depletion, many BG neurons inappropriately pattern their firing with respect to cortical slow oscillations (Magill et al., 2001; Belluscio et al., 2003; Walters et al., 2007). This might be relevant for the altered slow-wave sleep observed in people with PD (Zahed et al., 2021). However, it is unclear whether similar dysrhythmias arise in the Parkinsonian motor thalamus, hindering understanding of the wider functional impact of BG aberrations.

Studies of idiopathic PD suggest that beta-band dysrhythmia in the BG underpins bradykinesia/rigidity (Kühn et al., 2006, 2009; Ray et al., 2008; Sharott et al., 2014). Experiments in animals show the emergence of excessive beta rhythms throughout the dopamine-depleted BG is accompanied by abnormal increases or decreases in the average firing rates of constituent neurons (Mallet et al., 2008a, 2008b; Avila et al., 2010; Abdi et al., 2015; Sharott et al., 2017); these firing rate changes corroborate the predictions of the ‘direct/indirect pathways’ model of BG organization in Parkinsonism (Smith et al., 1998). This influential model further posits that, because GABAergic BG output neurons are hyperactive, motor thalamus neurons are hypoactive in Parkinsonism. However, reports suggest hyperactivity (Bosch-Bouju et al., 2014), hypoactivity (Schneider and Rothblat, 1996; Devergnas et al., 2016), or no change in firing rates in motor thalamus (Pessiglione et al., 2005; Anderson et al., 2015). One potential confounding factor is that motor thalamus is organized into discrete ‘input zones’, that is, a basal ganglia-recipient zone (BZ) and a cerebellar-recipient zone (CZ) (Nakamura et al., 2014). This important consideration aside, dopamine depletion alters other activity metrics in motor thalamus, including ‘burst firing’, pairwise correlations and oscillatory firing (Schneider and Rothblat, 1996; Pessiglione et al., 2005; Bosch-Bouju et al., 2014; Devergnas et al., 2016). Several of these alterations show that, like the BG, the motor thalamus is dysrhythmic in Parkinsonism. A single study of motor thalamus in dopamine-depleted rats suggests this might extend to beta-band dysrhythmia (Brazhnik et al., 2016). Critically, whether exaggerated beta oscillations are accompanied by changes in BZ neuron firing rates is unknown, as is the extent to which BZ neuronal ensembles might rhythmically synchronize their firing.

Elucidating the functional organization of the motor thalamus as a whole has benefited from direct comparisons of activity dynamics in the BZ and CZ, in both health and Parkinsonism (Anderson and Turner, 1991; Vitek et al., 1994; Guehl et al., 2003; Pessiglione et al., 2005; Ushimaru et al., 2012; Nakamura et al., 2014). The cerebellum might contribute to some compromised behaviors in PD (Wu and Hallett, 2013; Wichmann, 2019), and it is likely that CZ neuron activity is altered to some extent in Parkinsonism (Galvan et al., 2015). Because CZ is innervated by motor cortical regions exhibiting Parkinsonian beta oscillations (Mallet et al., 2008a, 2008b), dopamine depletion might also induce beta-band dysrhythmia in CZ (Basha et al., 2014).

To resolve these issues, we quantified the brain state-dependent firing of single cells and neuronal ensembles recorded in the anatomically-defined BZ and CZ of anesthetized dopamine-intact and dopamine-depleted rats. Our results emphasize that motor thalamus neurons are not hypoactive in Parkinsonism, but nevertheless engage in abnormal oscillatory activities in an input zone-selective manner.

## Materials and Methods

All experimental procedures were performed on adult male Sprague Dawley rats (Charles River) and were conducted in accordance with Animals (Scientific Procedures) Act, 1986 (United Kingdom).

### 6-Hydroxydopamine lesions of midbrain dopamine neurons

Unilateral 6-hydroxydopamine (6-OHDA) lesions were induced in 190–250 g rats, as previously detailed (Mallet et al., 2008a, 2008b; Abdi et al., 2015; Sharott et al., 2017). Briefly, the neurotoxin 6-OHDA (hydrochloride salt; Sigma) was dissolved in 0.9% w/v ice-cold NaCl solution containing 0.02% w/v ascorbate to a final concentration of 12 mg/ml. Approximately 25 min before the injection of 6-OHDA, all animals received desipramine (25 mg/kg, i.p.; Sigma) to minimize the uptake of 6-OHDA by noradrenergic neurons. Anesthesia was induced and maintained with 1.5–3% v/v isoflurane in O2, and animals were placed in a stereotaxic frame (Kopf). Body temperature was maintained at 37 ± 0.5 °C by a homeothermic heating device (Harvard Apparatus). Under stereotaxic control, 1 μl of 6-OHDA solution was injected near the medial forebrain bundle (4.1 mm posterior and 1.2–1.4 mm lateral of bregma, and 7.9 mm ventral to the dura (Paxinos and Watson, 2007)). Lesions were assessed 14–16 d after 6-OHDA injection by challenge with apomorphine (0.05 mg/kg, s.c.; Sigma) (Schwarting and Huston, 1996), and were considered successful when animals made ≥80 net contraversive rotations in 20 min (Sharott et al., 2017). Electrophysiological recordings (see below) were carried out in the thalamus or substantia nigra pars reticulata (SNr) ipsilateral to 6-OHDA lesions in anesthetized rats 21–51 d after surgery.

### In vivo electrophysiological recording and juxtacellular labeling of individual thalamic neurons

Recording and labeling experiments were performed in 36 anesthetized dopamine-intact rats (300–460 g) and 10 anesthetized 6-OHDA-lesioned rats (275–416 g at the time of recording), as previously described (Mallet et al., 2008a, 2008b, 2012; Nakamura et al., 2014; Abdi et al., 2015; Sharott et al., 2017). Briefly, anesthesia was induced with 4% v/v isoflurane in O2, and maintained with urethane (1.3 g/kg, i.p.; ethyl carbamate, Sigma), and supplemental doses of ketamine (30 mg/kg, i.p.; Willows Francis) and xylazine (3 mg/kg, i.p.; Bayer). Wound margins were infiltrated with local anesthetic (0.5% w/v bupivacaine, Astra). Animals were then placed in a stereotaxic frame (Kopf). Body temperature was maintained at 37 ± 0.5°C by a homeothermic heating device (Harvard Apparatus). Electrocorticograms (ECoGs) and respiration rate were monitored constantly to ensure the animals’ wellbeing. The epidural ECoG was recorded with a 1 mm-diameter screw above the frontal (somatic sensory-motor) cortex (4.2 mm rostral and 2.0 mm lateral of bregma (Paxinos and Watson, 2007)), and was referenced against a screw implanted above the ipsilateral cerebellum (Nakamura et al., 2014; Sharott et al., 2017). Raw ECoG was band-pass filtered (0.3–1500 Hz, −3 dB limits) and amplified (2000×; DPA-2FS filter/amplifier, NPI Electronic Instruments) before acquisition. Extracellular recordings of single-unit activity, that is, the action potentials (‘spikes’) fired by individual neurons, in the thalamus were made using standard-wall borosilicate glass electrodes (10–25 MΩ *in situ*; tip diameter 1.0–2.0 μm) containing 0.5 M NaCl solution and neurobiotin (1.5% w/v; Vector Laboratories, RRID:AB_2313575). Electrodes were lowered into the brain under stereotaxic guidance and using a computer-controlled stepper motor (IVM- 1000; Scientifica), which allowed electrode placements to be made with submicron precision. Electrode signals were amplified (10×) through the bridge circuitry of an Axoprobe-1A amplifier (Molecular Devices), AC-coupled, amplified another 100×, and filtered at 300–5000 Hz (DPA-2FS filter/amplifier). The ECoG and single-unit activity were each sampled at 17.9 kHz using a Power1401 Analog–Digital converter and a PC running Spike2 acquisition and analysis software (Cambridge Electronic Design). As described previously (Nakamura et al., 2014), single-unit activity in the thalamus was recorded during cortical slow-wave activity (SWA), which is similar to activity observed during natural sleep, and/or during episodes of spontaneous ‘cortical activation’, which contain patterns of activity that are more analogous to those observed during the awake, behaving state (Steriade, 2000). It is important to note that the neuronal activity patterns present under this anesthetic regime may only be qualitatively similar to those present in the unanesthetized brain. Nevertheless, the urethane-anesthetized animal still serves as a useful model for assessing the impact of extremes of brain state on functional connectivity within and between the basal ganglia, thalamus and cortex in dopamine-intact and Parkinsonian animals (Magill et al., 2006; Mallet et al., 2008a, 2008b; Sharott et al., 2012, 2017; Nakamura et al., 2014). Importantly, excessive beta oscillations arise (in a brain state-dependent manner) in the basal ganglia and motor cortex of 6-OHDA-lesioned rats under this anesthetic regimen (Mallet et al., 2008a, 2008b, 2012; Abdi et al., 2015; Sharott et al., 2017). Cortical activation was occasionally elicited by pinching a hindpaw for a few seconds. Note that we did not analyze neuronal activity recorded concurrently with the delivery of these sensory stimuli. Because the analyzed activity was recorded at least several minutes after the cessation of the brief pinch stimulus, it was also considered to be spontaneous (Mallet et al., 2008a; Nakamura et al., 2014; Sharott et al., 2017). The animals did not exhibit a marked change in respiration rate, and did not exhibit a hindpaw withdrawal reflex, in response to the pinch. Moreover, withdrawal reflexes were not present during episodes of prolonged cortical activation, thus indicating anesthesia was adequate throughout recordings.

Following electrophysiological recordings, some single thalamic neurons were juxtacellularly labeled with neurobiotin (Lacey et al., 2007; Nakamura et al., 2014). Briefly, positive current pulses (2–10 nA, 200 ms, 50% duty cycle) were applied until the single-unit activity became robustly entrained by the pulses. Single-unit entrainment resulted in just one neuron being labeled with neurobiotin. Two to six hours after labeling, animals were euthanized and transcardially perfused with 100 ml of 0.05 M PBS, pH 7.4 (PBS), followed by 300 ml of 4% w/v paraformaldehyde in 0.1 M phosphate buffer, pH 7.4 (PB). Brains were left overnight in fixative at 4°C, and then stored for 1–3 d in PBS at 4°C before sectioning (see below). Seventy nine of the thalamic neurons detailed herein were juxtacellularly labeled. These neurobiotin-labeled neurons were designated “identified” and precisely localized to different input zones of the motor thalamus (see below). The remaining (unlabeled) thalamic neurons (*n* = 100) were also included because, using stereotaxy and readouts from the stepper motor, we could accurately extrapolate their locations from those of identified neurons (recorded with the same glass electrodes in the same animals). Henceforth, we designate these unlabeled neurons as “extrapolated.” The identified and extrapolated thalamic neurons recorded in dopamine-intact rats with glass electrodes are those reported in Nakamura et al. (2014), but their firing properties have now been re-analyzed to address the issues underpinning the current study; specifically, we have performed new analysis of their firing with respect to ongoing cortical beta oscillations, and we have statistically compared thalamic neuron firing rates/patterns in dopamine-intact rats vs. 6-OHDA-lesioned rats.

### In vivo electrophysiological recording of thalamic and nigral activity with multielectrode arrays

Extracellular ‘wideband’ (0.1–6,000 Hz) recordings of neuronal activity were simultaneously made from numerous sites in the motor thalamus or SNr of urethane-anesthetized dopamine-intact rats (*n* = 7; 295–385 g) and 6-OHDA-lesioned rats (*n* = 11; 300–500 g at the time of recording) using linear electrode arrays with multiple, spatially-defined recording contacts (‘silicon probes’; A1×16-10mm-100-400 or A1×16-10mm-100-177, NeuroNexus), as previously described (Magill et al., 2006; Mallet et al., 2008b; Sharott et al., 2017). Each probe had 16 recording contacts arranged in a single vertical plane, with a contact separation of 100 μm. Depending on the probe used, each contact had an area of ∼400 μm^2^ (impedance of 0.9–1.2 MΩ, measured at 1000 Hz) or 177 μm^2^ (impedance of 1.7–2.0 MΩ). To enable *post hoc* histological verification of recording sites (see below), the backs of the silicon probes were evenly coated before each experiment with the red fluorescent dye 1,1’-dioctadecyl-3,3,3’,3’- tetramethylindocarbocyanine perchlorate (DiI; D3911, Invitrogen) by application of a 100 mg/ml DiI solution in acetone (Magill et al., 2006). The probe was manually advanced into the brain using a zero-drift micromanipulator (1760-61, Kopf) under stereotaxic control. The probes were cleaned after each experiment in a proteolytic enzyme solution (Magill et al., 2006). This was sufficient to ensure that contact impedances and recording performance were not altered by probe use and reuse. Monopolar probe signals were recorded using high-impedance unity-gain operational amplifiers (Advanced LinCMOS, Texas Instruments) and were referenced against a screw implanted above the contralateral cerebellum. After initial amplification, extracellular signals were further amplified (1000×) and low-pass filtered at 6000 Hz using programmable differential amplifiers (Lynx-8, Neuralynx). Electrocorticograms were also recorded as described above. The probe signals and ECoG were each sampled at 17.9 kHz using a Power1401 converter and a PC running Spike2 software. After the recording sessions, animals were euthanized and transcardially perfused with fixative (as described above) for *post hoc* histological analyses.

### Microinfusions of GABA into thalamus

For these experiments, urethane-anesthetized 6-OHDA-lesioned rats (*n* = 10; 300–500 g at the time of recording) were prepared for ECoG recordings as described above. A glass micropipette (tip diameter ∼ 30 µm) containing a 0.5M GABA solution (γ-aminobutyric acid; 0344, Tocris; dissolved in 0.9% w/v NaCl solution) was then advanced into the brain using a zero-drift micromanipulator under stereotaxic control. During sustained periods of cortical activation (>100 s of ensuing beta oscillations in ECoGs), 60 nl of the GABA solution was slowly infused (mean duration [± SEM]: 24.4 ± 1.0 s) into the motor thalamus using air pressure under manual control. Because the inactivation effects of such GABA microinfusions typically wear off within a few minutes (Kojima and Doupe, 2009), we were able to perform repeated infusions (using a minimal interval of 10 min) at one or more thalamic sites in a single animal; this also allowed us to negate the possibility of a spontaneous disappearance of cortical beta oscillations. To mark the microinfusion sites at the end of each experiment, 100–200 nl of a 0.9% w/v NaCl solution containing 0.04% w/v blue fluorescent microspheres (F8797, Invitrogen) was infused at the same stereotaxic coordinates through the same micropipette, as described previously (Nakamura and Morrison, 2007). Animals were then euthanized and transcardially perfused with fixative for *post hoc* histological analyses.

### Histology, immunofluorescence and microscopy

The fixed brains were cut into 50-μm thick sections in the parasagittal plane on a vibrating microtome (VT1000S; Leica Microsystems), collected in series, and washed in PBS. All the following reagent incubations were performed at room temperature.

To visualize neurobiotin-filled neurons, free-floating sections were washed in PBS and incubated overnight in Cy3-conjugated streptavidin (1:1000 dilution; PA43001, GE Healthcare) in “Triton-PBS” (PBS containing 0.3% v/v Triton X-100 [Sigma]). After washing, the sections were mounted on glass slides, coverslipped, and examined with an epifluorescence microscope (AxioPhot, Zeiss) to identify the neurobiotin-filled neurons. To delineate thalamic nuclei and map the location of each identified neuron (see Fig.1), sections containing the fluorescently-labeled somata were subsequently incubated overnight with a primary antibody mixture of rabbit anti-vesicular glutamate transporter 2 (VGluT2; 0.4 μg/mL of affinity-purified IgG; Hioki et al., 2003; a gift from Prof. Takeshi Kaneko, Kyoto University) and mouse anti-glutamic acid decarboxylase of 67 kDa (GAD67; 2 μg/mL; MAB5406, Millipore, RRID: AB_2278725) in Triton-PBS containing 1% v/v donkey serum (Jackson Immunoresearch; all the following antibody incubations were carried out with the same buffer). After washing with Triton-PBS, the sections were incubated for 2–4 h with a mixture of fluorophore-conjugated secondary antibodies (all raised in donkey): Anti-rabbit IgG (DyLight 649; 1:200; Jackson Immunoresearch) and anti-mouse IgG (Alexa Fluor 488; 1:200; Life Technologies). When necessary, the adjacent sections were incubated with NeuroTrace 500/525 (1:150; N-21480, Life Technologies), a green fluorescent Nissl stain, in Triton-PBS for 30 min to visualize cytoarchitecture. After washing, the fluorescently-labeled sections were mounted on glass slides, coverslipped, and examined with a laser-scanning confocal microscope (LSM710, Zeiss). To precisely localize the recorded neurons to distinct thalamic nuclei, fluorescent images of the thalamus around the neurobiotin-filled neurons were taken at a low magnification with a 5× objective lens (EC Plan-Neofluar, numerical aperture 0.16; Zeiss), a pinhole thoroughly opened (i.e. in “non-confocal” mode), and a zoom factor of 0.6. Appropriate sets of laser beams and emission windows were used for Alexa Fluor 488 (excitation 488 nm, emission 492–544 nm), Cy3 (excitation 543 nm, emission 552–639 nm), and DyLight 649 (excitation 633 nm, emission 639–757 nm). Images of each of the channels were taken separately and sequentially to negate possible “bleed through” of signal across channels. Images were combined into montages and, when necessary, images from the adjacent sections were overlaid and aligned using graphic software (Canvas 12, ACD Systems, RRID:SCR_014288). The two input zones of the motor thalamus (BZ and CZ) were delineated on the basis of their distinctive distributions of VGluT2 and GAD67 immunoreactivities (Kuramoto et al., 2009, 2011, 2015; Nakamura et al., 2014). Only identified neurons located >50 μm away from the borders of BZ or CZ were analyzed. Extrapolated neurons had to be located >100 μm away from these borders to be included in the analyses. The dendrites of thalamocortical neurons in the rat motor thalamus only rarely radiate >200 μm from the parent somata (Kuramoto et al., 2009, 2015). Thus, most of the proximal dendrites of our neurons were likely to be confined to just one zone.

**Figure 1.**
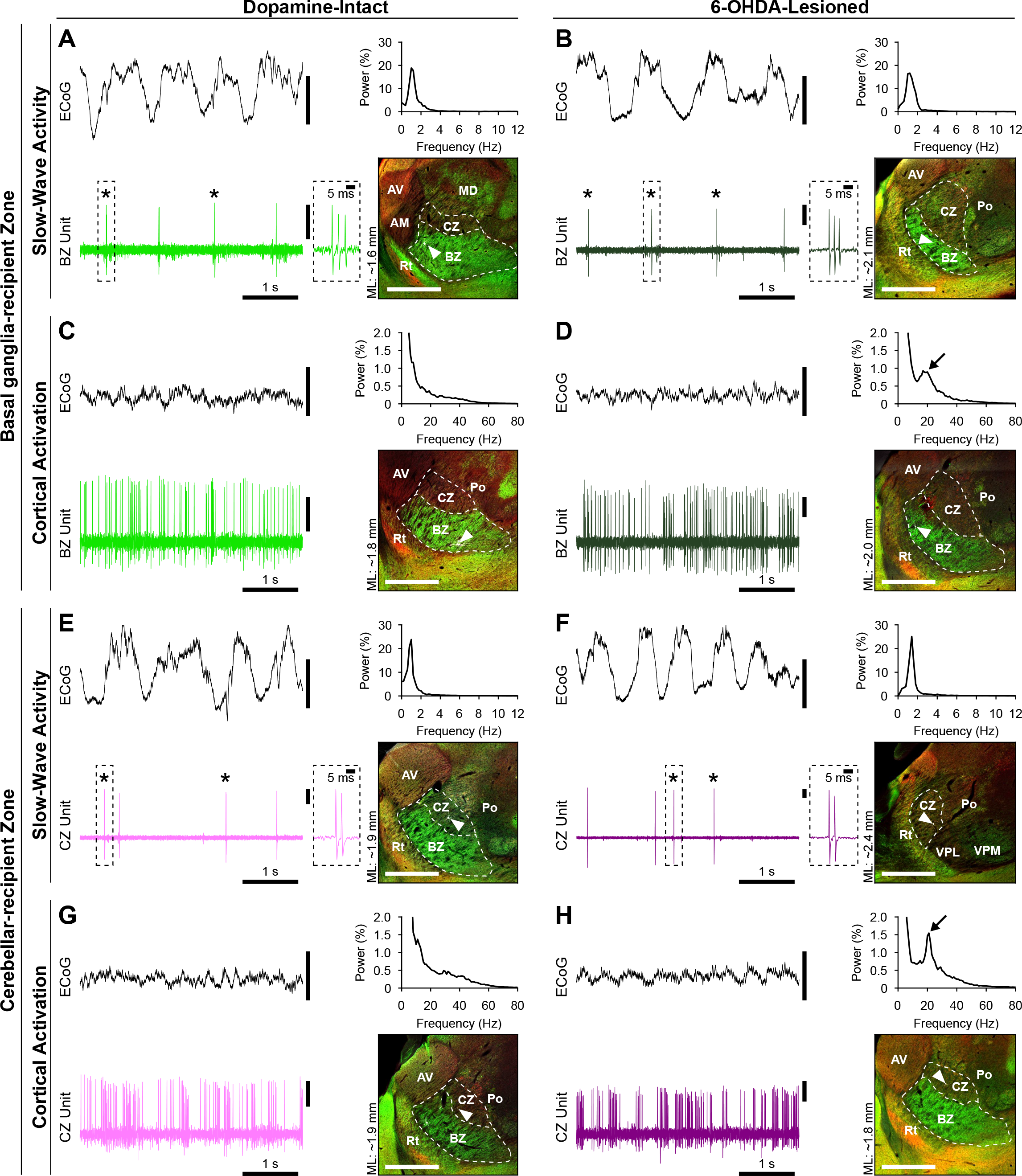
Brain state-dependent firing of identified neurons in the basal ganglia- recipient zone and cerebellar-recipient zone of the motor thalamus in dopamine-intact and 6-OHDA-lesioned rats. ***A***, *Left,* Spontaneous firing of a thalamic unit during cortical slow-wave activity in a dopamine-intact rat. Brain state was defined according to simultaneous electrocorticogram (ECoG) recordings; during SWA, the ECoG was dominated by slow (∼1 Hz) oscillations, as further verified in the ECoG power spectrum (*upper right*). The thalamic unit fired stereotypical low-threshold Ca^2+^ spike bursts (indicated with asterisks) on most cycles of the cortical slow oscillation. Such bursts were characterized by 2–6 action potentials fired in rapid succession, often with a progressive attenuation of action potential amplitude during the burst (a typical burst is highlighted by the dashed box and also shown at higher temporal resolution in the *inset*). *Right*, Subsequent to electrophysiological recording with a glass electrode, the same neuron was juxtacellularly filled with neurobiotin, fluorescently labeled, and identified (white, arrowhead). The neuron was then localized to the basal ganglia-recipient zone (BZ) of motor thalamus, which was demarcated by intense GAD67 immunoreactivity (green) and sparse VGluT2 immunoreactivity (red) in the parasagittal tissue section (∼1.6 mm lateral of bregma [ML]). ***B***, Spontaneous firing of an identified BZ neuron during SWA in a 6-OHDA-lesioned rat. ***C***, Spontaneous firing of an identified BZ neuron recorded during cortical activation, as verified by a relative paucity of ECoG slow oscillations, in a dopamine-intact rat. Note that the BZ neuron does not fire low-threshold spike bursts during cortical activation. ***D***, Spontaneous firing of an identified BZ neuron during cortical activation in a lesioned rat. Note the distinct peak at beta frequencies (black arrow) in the ECoG power spectrum. ***E***, Recording made during cortical SWA in a dopamine-intact rat of a neuron that was subsequently localized to the cerebellar-recipient zone (CZ) of motor thalamus, as delineated by sparse GAD67 and moderate VGluT2 immunoreactivities. ***F***, Spontaneous firing of an identified CZ neuron during SWA in a lesioned rat. ***G***, Spontaneous firing of an identified CZ neuron during cortical activation in a dopamine-intact rat. ***H***, Spontaneous firing of an identified CZ neuron during cortical activation in a lesioned rat. In all parasagittal sections, rostral is towards the left, and dorsal is towards the top, with the borders of BZ and CZ indicated with dashed white lines. AM, anteromedial thalamic nucleus; AV, anteroventral thalamic nucleus; MD, mediodorsal thalamic nucleus; Po, posterior nuclear group; Rt, thalamic reticular nucleus; VPL, ventral posterolateral thalamic nucleus; VPM, ventral posteromedial thalamic nucleus. Scale bars in fluorescence images: 1 mm. Vertical calibration bars: 0.5 mV.

To determine the locations of silicon probe recording sites (see Figs.6 and 8), tissue sections were mounted on glass slides in PBS, coverslipped, and examined with a microscope capable of fluorescent and brightfield imaging (Axio Imager.M2, Zeiss). In those sections containing DiI signal, images of fluorescence (43 Cy3 filter) and transmitted light were then taken with a 1.25× objective (EC Plan-Neofluar, numerical aperture 0.03; Zeiss) to record the probe penetration tracks. In many cases, the best quality DiI images were obtained at this stage, because the subsequent immunofluorescence protocol tended to ‘wash out’ the DiI. At the start of the immunofluorescence protocol, the DiI-containing sections were unmounted and then heat treated as a means of antigen retrieval (80°C for 30 min in 10 mM citrate-NaOH buffer, pH6.0). After washing with PBS, the sections were incubated overnight with a primary antibody mixture of guinea pig anti-glycine transporter 2 (GlyT2; 1:10,000; AB1773, Merck, RRID:AB_90953) and mouse anti-GAD67 (1 µg/ml; MAB5406, Millipore, RRID:AB_2278725) for the thalamus, or mouse anti-GAD67 antibody alone for the SNr, in Triton-PBS containing 10% v/v donkey serum (Jackson Immunoresearch; all the following antibody incubations were carried out with the same buffer). After washing with Triton-PBS, the sections were incubated for 2–4 h with a mixture of fluorophore-conjugated secondary antibodies (all raised in donkey, 1:500, Jackson Immunoresearch): Anti-guinea pig IgG (DyLight 488) and anti-mouse IgG (Alexa Fluor 647) for the thalamus, or anti-mouse IgG alone for the SNr. A laser-scanning confocal microscope was used to take low-magnification fluorescent images of the thalamus around the DiI signal, as described above. To correct for tissue shrinkage (<10%) that occurred during the immunofluorescence protocol, images taken after immunofluorescence were scaled and aligned with images taken before immunofluorescence, using Canvas 12 software. Judging from the known stereotaxic distances between probe penetrations in a single plane (see Figs.6*A,A’* and *8A,A’*), we estimated the difference between the scaling in the tissue *in vivo* during recordings and the scaling in the images before immunofluorescence to be <10%. The most ventral DiI deposit in each penetration track was considered to be the location of the probe tip; extrapolating from this, the estimated positions of the probe recording contacts were plotted on the images of DiI signal and immunofluorescence (see Figs.6*A,A’* and *8A,A’*). Thalamic nuclei/zones and the SNr were identified according to immunofluorescence images and each recording contact was assigned a location tag for group analyses. When the estimated locations of probe contacts were on or close to (≤50 µm) the borders between structures, those recording sites were tagged as “border” and were excluded from group analyses (see Fig. 6*B*–*D*, 8*B*–*D*).

To determine the sites of GABA microinfusion, we used an anatomical analysis pipeline similar to that for localizing silicon probes, with the key difference being that glass pipette tracks were visualized with blue fluorescent microspheres (see Fig.10). Thalamic nuclei/zones were identified with immunofluorescence for GlyT2, VGluT2 and GAD67, using the primary antibodies described above and revealed with secondary antibodies that were respectively conjugated to DyLight 488, Cy3, and Alexa Fluor 647. A laser-scanning confocal microscope was first used to take low-magnification fluorescent images of the blue microspheres (excitation 405 nm, emission 409–559 nm) in the tissue. Using the same objective and zoom, images of DyLight 488, Cy3, and Alexa Fluor 647 signals were then taken sequentially and separately (as above) to map thalamic structures. After registration of images, the ends of the pipette tracks (considered to be the locations of the pipette tips) were assigned a location tag.

### Analysis of ECoGs and basic firing parameters of single units

Electrocorticogram data from each recording session were visually inspected and epochs of robust cortical SWA or cortical activation were selected according to the previously described characteristics of these brain states (Mallet et al., 2008b; Nakamura et al., 2014; Sharott et al., 2017). A 100 s portion of the glass electrode or silicon probe data concomitantly recorded during each defined brain state was isolated and used for statistical analyses. Silicon probe data were high-pass filtered off-line at 300 Hz (Spike2, finite impulse response filter) to isolate unit activity. Putative single-unit activity was isolated with standard “spike sorting” procedures (Mallet et al., 2008a), including template matching, principal component analysis, and supervised clustering (Spike2). Isolation of a single unit was verified by the presence of a distinct refractory period in the interspike interval (ISI) histogram. Only neurons in which <1% of all ISIs were <2 ms were analyzed in this study. Single-unit activity was converted so that each spike was represented by a single digital event (Spike2). The recorded signals and sorted spike trains were then resampled at 17 kHz with Spike2 and exported to MATLAB (MathWorks, RRID:SCR_001622) for further analysis. Spike trains were assumed to be realizations of stationary stochastic point processes. The mean firing rate (spikes/s) of individual neurons was calculated from the total number of spikes per 100 s data epoch. Variability of firing was assessed using a metric related to the coefficient of variation (CV) of the ISI, the mean CV2 (Holt et al., 1996); the lower the CV2 value, the more regular the unit activity.

### Detection of low-threshold Ca2^+^ spike bursts fired by thalamic neurons

Low-threshold Ca^2+^ spike (LTS) bursts were classified as such using custom Spike2 scripts, according to previously defined criteria for identifying the LTS bursts in extracellular unit recordings (Lacey et al., 2007; Nakamura et al., 2014): 1) At least 2 action potentials with an ISI of ≤5 ms but with a preceding silent period of >100 ms (Lu et al., 1992); and 2) a maximum ISI of 10 ms was used to define the end of a LTS burst (Fanselow et al., 2001).

### Detection of the burst firing of SNr neurons

Analyses of SNr neuron burst firing in lesioned rats was carried out on single units that displayed high variability in their ISIs (CV > 1.0) during SWA. To detect the onsets and offsets of burst firing, SNr spike trains (*n* = 14, from 5 rats) were first converted into a spike density function (SDF; see Fig.4*G*) (Szucs, 1998). The point process data were thus converted to continuous time series (changes in firing probability over time) by convolution (with MATLAB function conv) of a Gaussian kernel with a sigma of 0.051 s (selected so that 95% of spikes are within the range of [−3σ, 3σ]), as prepared by MATLAB function normpdf. Peaks and troughs in the SDF waveform were then detected using MATLAB function findpeaks, using a minimal peak/trough duration of 2σ and, for peak detection only, setting the minimal peak height to be the maxima of the Gaussian kernel. A threshold of the SDF waveform was then set as the mean of the median peaks and median troughs (Fig.4*G*). Within the epochs that the SDF waveform was equal to or higher than this threshold, the spike events that were closest to the times that the SDF rose above or fell below the threshold were considered to be the onsets and offsets of bursts, respectively.

### Analysis of phase-locked firing of single units, including circular statistics

To investigate how the activity of individual thalamic and nigral neurons varied in time with respect to ongoing cortical network activity, we analyzed the instantaneous phase relationships between thalamic/nigral spike times and cortical oscillations in specific frequency bands (Sharott et al., 2012, 2017; Nakamura et al., 2014; Garas et al., 2016). Signal analyses were performed using MATLAB. Electrocorticogram signals containing robust SWA or cortical activation were initially downsampled to 1024 Hz (MATLAB function resample) and then digitally band-pass filtered to isolate slow (0.4–1.6 Hz) or beta (15–30 Hz) oscillations, respectively (second- and fifth-order zero-phase Butterworth filters for slow and beta oscillations, respectively, using MATLAB function filtfilt). Subsequently, the instantaneous phase and power of the ECoG in these frequency bands were separately calculated from the analytic signal obtained via the Hilbert transform (Lachaux et al., 1999). In this formalism, peaks in the ECoG oscillations correspond to a phase of 0° and troughs to a phase of 180°. Linear phase histograms, circular phase plots, and circular statistical measures were calculated using the instantaneous phase values for each spike. Descriptive and inferential circular statistics were then calculated using the CircStat toolbox (Berens, 2009) for MATLAB. The phase-locked firing of each neuron with respect to cortical oscillations was represented by a vector whose angle and length (bound between 0 to 1; the closer to 1, the more concentrated the angles) were respectively defined as the circular mean (circ_mean of CircStat) and the mean resultant vector length (simply referred to as ‘vector length’; circ_r of CirtStat) of the instantaneous phase values of spikes. For the calculation of vector lengths and statistical comparisons, we included only those neurons that fired ≥40 spikes during the entire analyzed epoch (100 s). These ‘qualifying’ neurons were then tested for significantly phase locked firing (defined as having *p* < 0.05 in Rayleigh’s Uniformity Test; circ_rtest of CircStat). The null hypothesis for Rayleigh’s test was that the spike data were distributed in a uniform manner across/throughout phase. We and others have previously remarked that the non-sinusoidal nature of some field potential oscillations, such as the cortical slow oscillation, can confound standard circular statistics, especially Rayleigh’s test (Siapas et al., 2005; Mallet et al., 2008a, 2008b; Sharott et al., 2012; Nakamura et al., 2014). Thus, for analysis of neuron firing relationships with cortical slow oscillations, Rayleigh’s tests were only carried out after any phase non-uniformities of the slow oscillations were corrected with the empirical cumulative distribution function (MATLAB ecdf) (Siapas et al., 2005; Nakamura et al., 2014; Abdi et al., 2015; Garas et al., 2016). For each of the neurons that were significantly phase-locked using these criteria, the mean phase angle was calculated. Because some datasets did not meet the requirements for parametric testing, differences in the median phase angles of groups of neurons were tested using the non-parametric common median test (circ_cmtest of CircStat) (Fisher, 1995). The null hypothesis for the common median test is that all tested groups have the same circular median. The vector length was used to quantify the level of phase locking around the mean phase for individual neurons (computed using the angles of each spike) and for populations of neurons (computed using the mean phase for each neuron). Where single-unit data are displayed in circular plots, lines radiating from the center are the vectors of the preferred phases of firing (with the center and perimeter of the outer grid circle representing vector lengths of 0 and 1, respectively); thin lines indicate preferred firing of individual neurons, whereas thick lines indicate population vectors. The small open circles on the perimeter represent the preferred phases of each neuron.

### Extraction and conditioning of background-unit activity signals for time series analyses

We analyzed ‘background-unit activity’ (BUA) signals recorded with silicon probes, a representation of the summed firing of small, local neuronal populations that is conceptually distinct from multiunit activity and LFPs (Moran and Bar-Gad, 2010). These BUA signals were isolated from the wideband recordings made with the probes by initial high-pass filtering off-line at 300 Hz (Spike2, finite impulse response filter) and the removal of any large-amplitude action potentials that could potentially distort the signals and bias analyses, as per Moran et al (Moran et al., 2008) (Fig.6-1*A*–*F*). Large-amplitude action potentials were defined as those exceeding 3 standard deviations of the entire high-pass filtered signal, and data points around these large action potentials were removed and replaced with another randomly-selected part of the recording that did not contain similarly large action potentials (MATLAB createBUA function, a gift from Dr Izhar Bar-Gad). The windows for spike removal were generally set to −1.5 and +2.0 ms before and after the peak of the large action potentials, with a few exceptions (2 and 14 out of 673 channels for before and after a spike, respectively) for which wider (up to −3.0 and +3.0 ms) windows were used to avoid artifacts as assessed by visual inspection. The BUA signals were then full-width rectified (Journée, 1983; Myers et al., 2003; Moran et al., 2008) and mean subtracted to remove the DC component that was created by rectification (Fig.6-1*G*–*I*). Finally, the signals were low-pass filtered at 300 Hz (zero-phase shift Butterworth filter with the order of 3) and downsampled to 1024 Hz (Fig.6-1*J*–*L*) before further time series analyses.

### Analysis of phase-locked BUA signals, including circular statistics

To investigate how thalamic and nigral BUA varied in time with respect to ongoing cortical beta oscillations, we analyzed ‘phase-averaged waveforms’ of BUA signals, that is, the average of BUA voltage (µV) in each bin (size = 5°) of the instantaneous phase values of the ECoG band-pass filtered at 15–30 Hz (zero-phase shift Butterworth filter with the order of 3). A sample vector 𝑉 for an individual BUA signal is defined in the complex plane as the double of an average of complex number-based vector representations of the instantaneous phase values and the values of BUA signals of each data point, *i.e. :* 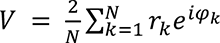, where 𝜑 _𝑘_ and 𝑟 _𝑘_ represent the instantaneous phase in radians and the value of BUA signal (signed) in µV for the 𝑘th data point, respectively, 𝑁 is the number of data points, and 𝑖 is the imaginary unit. The average was doubled to reflect the amplitude difference between the positive and negative deflections. If the phase-averaged waveform of a signal is an ideal sinusoidal curve, the sample vector length |𝑉| is identical to the peak-to-peak amplitude in µV.

In order to assess the non-uniformity (i.e. significant phase modulation) of BUA signals in relation to cortical beta oscillations (Figs.7*G,H* and 9*G,H*), data points were circularly shifted in a random manner within each cycle of the beta oscillations (MATLAB circshift). This maintained the waveform of BUA signals within a cycle while randomizing their phase relationships to ECoG. A similar approach has been used elsewhere (von Nicolai et al., 2014). We randomly chose 10 BUA signals in BZ or SNr and performed 1000 iterations of random shifting of each signal to generate a histogram based on an empirical cumulative distribution function of vector lengths of the shifted data (MATLAB ecdfhist). A BUA signal was considered to be ‘significantly modulated’ in relation to ECoG beta oscillations when the sample vector length was longer than 99.9% of the sample vector lengths of the shifted data. Where significantly-modulated BUA signals are displayed in circular plots (Figs.7*I* and 9*I*), lines radiating from the center are the vectors of their preferred phases (with the center and perimeter of the outer grid circle representing vector lengths of 0 and 1.2 µV, respectively); thin lines indicate the preferred phase of individual BUA signals, whereas thick lines indicate population vectors. The small open circles on the perimeter represent the preferred phases of each BUA signal. Group analysis of the sample vectors for BUA signals was carried out as for group analysis of single-unit data (see above).

### Spectral analyses

Electrocorticograms were downsampled to 1024 Hz before spectral analyses. Spectral parameters for ECoG and BUA time series were evaluated using Fast Fourier Transform (FFT). Power and coherence spectra were calculated with MATLAB pwelch and mscohere functions, with a FFT size of 5120 for signals recorded during SWA (giving a frequency resolution of 0.2 Hz) and a FFT size of 1024 for signals recorded cortical activation (1.0 Hz resolution). The overlap of FFT windows was 50%. For power spectra, each individual power spectrum was normalized to give “% relative power” unless otherwise stated. This was achieved by calculating the spectral power in each frequency bin as a percentage of the total power between 0.4 and 100 Hz (for analysis of SWA) or between 1 and 100 Hz (for analysis of cortical activation). For statistical comparisons, the sum of power or the average of coherence across all frequency bins in the band of interest (i.e. 0.4–1.6 Hz or 15–30 Hz) was calculated, giving a single value for each recording.

### Analysis of effects of GABA microinfusion

We characterized the extent to which microinfusions of GABA into thalamus influenced ongoing cortical beta oscillations. To visualize changes in ECoG power at beta frequencies (15–30 Hz) over time and across experiments, we plotted beta-band power normalized to that in a ‘pre-GABA period’ set as the 100 s immediately before GABA infusion onset. The effect size (%) of GABA infusion was defined as (1 − 𝑥) × 100 [%], where 𝑥 is the ratio of the total ECoG beta-band power during the 50–150 s after GABA infusion onset to that during the pre-GABA period.

### Statistical analyses

For each experiment, descriptions of critical variables (e.g., number of animals, neurons, and other samples evaluated) as well as statistical design can be found in the Results. The Shapiro–Wilk test (swtest by Ahmed BenSaïda), was used to judge whether noncircular datasets were normally distributed (*p* < 0.05 to reject). Because some data sets were not normally distributed, we used nonparametric statistical testing for these data throughout. The Mann–Whitney U test (MWUT; MATLAB ranksum) was used for comparisons of unpaired data. For multiple group comparisons, we performed a Kruskal–Wallis ANOVA on ranks (MATLAB kruskalwallis), with Dunn’s test (MATLAB multcompare) for further *post hoc* definition of comparisons. Significance for all statistical tests was set at *p* < 0.05 (exact *p* values are given in the text). Data are represented as group means ± SEMs unless stated otherwise. All box plots in figures show the individual samples (circles), medians, the interquartile ranges (box), and the non-outlier values closest to the first and third quartiles (whiskers).

## Results

The overall aim of this study was to define how the chronic depletion of dopamine, as occurs in PD, alters the spatial and temporal organization of electrical activity within the two major input zones of the motor thalamus *in vivo*. Emphasis was placed on defining the extent to which the action potential firing of BZ and CZ neurons becomes dysrhythmic during, and with respect to, the slow oscillations (0.4–1.6 Hz) and beta oscillations (15–30 Hz) emerging in cortico-basal ganglia circuits in a brain state- and dopamine-dependent manner. To address this, we first recorded individual, identified neurons in the BZ and CZ of anesthetized dopamine-intact rats and dopamine-depleted (6-OHDA-lesioned) rats during two well-defined and controlled brain states, slow-wave activity (SWA) and cortical activation. To gain further insights into the activity dynamics of larger neuronal populations during cortical activation, we sampled background-unit activities from numerous sites in and around the BZ and CZ using linear multi-electrode arrays. Further functional context was provided by recording the activities of neurons in the substantia nigra pars reticulata (SNr), one of the BG output nuclei that targets BZ, as well as by examining the effects of pharmacological perturbations of the BZ and CZ.

### Dopamine depletion does not decrease the firing rates of neurons in the motor thalamus during cortical slow-wave activity or cortical activation

The influential direct/indirect pathways model of BG organization in Parkinsonism predicts that motor thalamus neurons are hypoactive in Parkinsonism (DeLong, 1990; Smith et al., 1998). Ongoing brain state provides critical context when testing the validity of the model’s predictions; not only are the firing rates of motor thalamus neurons in dopamine-intact rodents exquisitely dependent on brain state (Ushimaru et al., 2012; Nakamura et al., 2014), but it has also been established that dopamine depletion only alters the firing rates of BG neurons in the expected manner during certain brain states (Abdi et al., 2015; Sharott et al., 2017; Kovaleski et al., 2020). With this in mind, and throughout this study, we interrogated neuronal activity dynamics in the motor thalamus in the context of cortical SWA and cortical activation, as verified in simultaneous recordings of ipsilateral frontal electrocorticograms (Magill et al., 2006; Mallet et al., 2008a, 2008b; Nakamura et al., 2014).

Using glass electrodes, we recorded the spontaneous action potential discharges (spikes) of 137 single units (neurons) in the motor thalamus of dopamine-intact control rats (*n* = 36), and 42 neurons in the motor thalamus of 6-OHDA-lesioned rats (*n* = 10), during cortical SWA and/or cortical activation. Of these neurons, 46% (63 of 137) and 38% (16 of 42) were unequivocally identified, that is, after electrophysiological characterization, they were juxtacellularly labeled with neurobiotin, and their somata were precisely localized to the BZ or CZ (Fig.1). As described previously (Kuramoto et al., 2009, 2011; Bosch-Bouju et al., 2014; Nakamura et al., 2014), we used markers of specific groups of GABAergic axon terminals or glutamatergic axon terminals (*i.e.* GAD67 and VGluT2, respectively) to define the boundaries of the BZ and CZ (Fig.1). During periods of robust SWA in ipsilateral frontal cortex, the activity of identified BZ neurons in both dopamine-intact rats and lesioned rats was typified by a relatively low mean firing rate (<4 spikes/s) and a propensity to fire discrete bursts of spikes (Fig.1*A*,*B*). These bursts were exemplified by 2–6 spikes fired in rapid succession (instantaneous intraburst rates of >150 spikes/s), with a progressive decrease in spike amplitude (Fig.1*A*,*B* *insets*). Many of these bursts satisfied the criteria for stereotypical low-threshold Ca^2+^ spike (LTS) bursts (see Materials and Methods). Moreover, spikes were often fired in time with cortical slow (∼1 Hz) oscillations (Fig.1*A*,*B*). The activities of neurons in dopamine-intact and lesioned rats showed clear brain state-dependency. Thus, during cortical activation, which was exemplified by a relative paucity of cortical slow oscillations (and, in the case of lesioned rats only, the emergence of exaggerated beta oscillations), the activity of identified BZ neurons in both dopamine-intact and lesioned rats was typified by relatively high mean firing rates (>10 spikes/s) and a “tonic” irregular firing pattern (Fig.1*C*,*D*). Neurons in BZ seldom fired LTS bursts during cortical activation. Importantly, the activity of identified CZ neurons was qualitatively similar to that of BZ neurons, irrespective of brain state and whether recordings were made in dopamine-intact rats and or in lesioned rats (Fig.1*E*–*H*).

We next quantitatively assessed whether dopamine depletion altered the basic firing properties of BZ neurons or CZ neurons. For these analyses, we pooled together all identified and “extrapolated” neurons, that is, the unlabeled neurons whose locations could be accurately extrapolated from those of identified neurons (see Materials and Methods). We first considered thalamic activity recorded during cortical SWA (Fig.2*A*–*F*). On average, the firing rates of BZ neurons in lesioned rats (2.76 ± 0.22 spikes/s [mean ± SEM]; *n* = 13 neurons) were slightly, but significantly, higher (*p* = 0.002, MWUT; Fig.2*A*) than those of BZ neurons in dopamine-intact rats (1.88 ± 0.06 spikes/s; *n* = 48 neurons). In contrast, the firing rates of CZ neurons in lesioned rats (2.51 ± 0.41 spikes/s; *n* = 14 neurons) were similar (*p* = 0.149, MWUT; Fig.2*B*) to those of CZ neurons in dopamine-intact rats (1.93 ± 0.09 spikes/s; *n* = 80 neurons). Dopamine depletion did not alter the firing variability (as indexed by CV2 measures) of either BZ neurons (*p* = 0.679, MWUT; Fig.2*C*) or CZ neurons (*p* = 0.061, MWUT; Fig.2*D*). During SWA, all motor thalamus neurons fired numerous LTS bursts (Fig.1); on average, ∼75% of all spikes fired by BZ and CZ neurons during SWA occurred within LTS bursts (Fig.2*E*,*F*). Dopamine depletion did not alter the propensities of BZ and CZ neurons to fire spikes in LTS bursts (*p* = 0.465 and 0.342 for BZ and CZ neurons, respectively, MWUT; Fig.2*E*,*F*). We then considered thalamic activity recorded during cortical activation (Fig.2*G*–*L*). On average, the firing rates of BZ neurons in lesioned rats (20.48 ± 2.66 spikes/s; *n* = 11 neurons) were similar (*p* = 0.237, MWUT; Fig.2*G*) to those of BZ neurons in dopamine-intact rats (16.49 ± 0.52 spikes/s; *n* = 32 neurons). In contrast, the firing rates of CZ neurons in lesioned rats (24.01 ± 2.29 spikes/s; *n* = 8 neurons) were significantly higher (*p* = 0.013, MWUT; Fig.2*H*) than those of CZ neurons in control rats (16.39 ± 0.70 spikes/s; *n* = 30 neurons). Dopamine depletion increased the firing variability of BZ neurons (*p* = 0.006, MWUT; Fig.2*I*), but did not alter the firing variability of CZ neurons (*p* = 0.508, MWUT; Fig.2*J*). During cortical activation, the proportions of all spikes included in LTS bursts were very low for neurons in both input zones (0.23 ± 0.15% and 0.07 ± 0.02% for BZ neurons in lesioned and dopamine-intact rats, respectively; 0.10 ± 0.08% and 0.07 ± 0.03% for CZ neurons in lesioned and dopamine-intact rats; Fig.2*K*,*L*). Dopamine depletion did not alter the propensities of BZ and CZ neurons to fire LTS bursts during cortical activation (*p* = 0.171 and 0.543 for BZ and CZ neurons, respectively, MWUT; Fig.2*K*,*L*).

**Figure 2.**
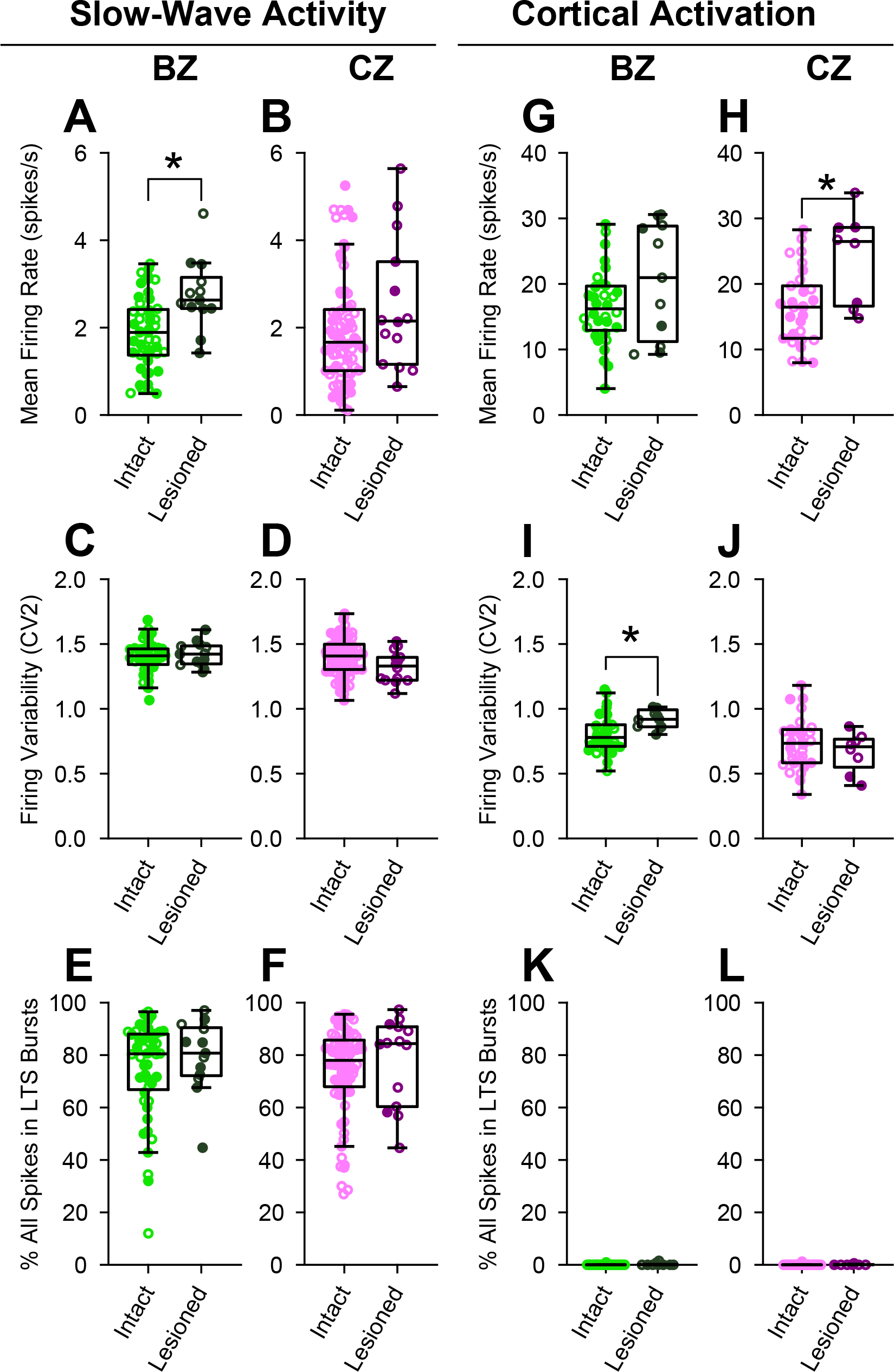
Quantitative comparisons of the firing rates and patterns of BZ and CZ neurons during two brain states in dopamine-intact and 6-OHDA-lesioned rats. Mean firing rates (***A***, ***B***, ***G*** and ***H***), firing variability, as indexed by the coefficients of variation (CV2) of interspike intervals (***C***, ***D***, ***I*** and ***J***), and mean percentage of all spikes occurring in low-threshold Ca^2+^ spike bursts (***E***, ***F***, ***K*** and ***L***) for all individual neurons recorded with glass electrodes in the BZ (light/dark greens) and in the CZ (pink/purple) of the motor thalamus. ***A–F***, Thalamic activity parameters during SWA. Dopamine depletion was associated with an increase in the mean firing rates of BZ neurons (only), but did not alter the firing variabilities of either BZ or CZ neurons during SWA. ***G–L***, Thalamic activity parameters during cortical activation. On average, the firing rates of CZ neurons, but not BZ neurons, were increased in lesioned rats. The activity patterns of motor thalamic neurons were intimately related to brain state; during cortical activation, their firing rates were relatively high and they rarely fired LTS bursts. Box plots in this and subsequent figures denote the non-outlier values closest to the first and third quartiles (whiskers), interquartile range, and medians. Data from individual identified and extrapolated neurons are shown as filled and open circles, respectively. \**p* < 0.05 (exact *p* values are in Results text), Mann–Whitney U tests.

In summary, these recordings of individual neurons accurately localized to one or the other input zone of the motor thalamus show that the firing rates of BZ neurons are not abnormally decreased after chronic dopamine depletion. These data also suggest that, when carefully controlling for two brain states, dopamine depletion does not increase the prevalence of LTS burst firing in either the BZ or CZ.

### The firing of BZ neurons, but not CZ neurons, is aberrantly phase-locked to cortical slow oscillations after dopamine depletion

Dopamine depletion can alter the patterning of BG neuron discharges during, and with respect to, the stereotyped cortical slow oscillation (Magill et al., 2001; Mallet et al., 2006; Walters et al., 2007; Zold et al., 2012; Abdi et al., 2015). We addressed whether this also holds true for motor thalamus neurons. During SWA, the normalized power of ECoG slow oscillations (defined as 0.4–1.6 Hz Nakamura et al., 2014; Sharott et al., 2017) simultaneously recorded with BZ neurons was on average similar between dopamine-intact rats and lesioned rats (*p* = 0.052, MWUT; Fig.3*A*). The power in ECoGs recorded with CZ neurons was also similar across animal groups (*p* = 0.920, MWUT; Fig.3*A*). The power of slow oscillations in single-unit activities in BZ was on average slightly higher in dopamine-intact rats than in lesioned rats (*p* = 0.035, MWUT; Fig.3*B*), as was the coherence between ECoGs and BZ units at slow oscillation frequencies (*p* = 0.046, MWUT; Fig.3*C*). The equivalent metrics in CZ were unaffected by dopamine depletion (*p* = 0.441 for CZ units, MWUT; Fig.3*B*; *p* = 0.241 for ECoG– CZ coherence, MWUT; Fig.3*C*). These results suggest that dopamine depletion selectively alters the relationship between activities in frontal cortex and BZ. To test this further, we used the Hilbert transform to analyze the instantaneous phase of the spiking of thalamic neurons with respect to cortical slow oscillations at 0.4–1.6 Hz (Nakamura et al., 2014; Abdi et al., 2015; Garas et al., 2016; Sharott et al., 2017). On average, BZ neurons in dopamine-intact rats preferentially fired at the ascending phase of cortical slow oscillations (Fig.3*D*), that is, at around the phase that cortical neurons transition from silence to coordinated firing (Sakata and Harris, 2009; Chauvette et al., 2010). This finely-timed firing of BZ neurons was also evident in their individual phase histograms (Fig.3*E*). Circular statistical analyses revealed that the spikes of all BZ neurons (*n* = 48) were significantly phase-locked to the slow oscillations (*p* < 0.05, Rayleigh’s Uniformity Test). Circular plots of the preferred phases of these phase-locked BZ neurons also demonstrated their strong tendency to fire at the ascending phase of the slow oscillations (Fig.3*F*); the mean angle of the preferred firing phases for the BZ neuron group was 283.4 ± 3.8° (Fig.3*J*). Dopamine depletion profoundly disturbed the temporal coupling (phase locking) of BZ neuron firing to cortical slow oscillations (Fig.3*D*–*F*,*J*). On average, and when compared to BZ neurons in dopamine-intact rats, the peak of activity of BZ neurons in lesioned rats appeared smaller and broader in the linear phase histogram, indicating weaker and more variable phase locking as a population after dopamine depletion (Fig.3*D*). Furthermore, most BZ neurons in lesioned rats preferentially fired at the descending phase of cortical slow oscillations (Fig.3*D–F*), with a mean angle of firing of 31.7 ± 15.4° for the BZ neuron group (Fig.3*J*). Accordingly, dopamine depletion resulted in a large (∼100°) and significant (*p* = 0.001, common median test) shift in the mean angles of firing of phase-locked BZ neurons. In stark contrast, dopamine depletion did not alter the temporal coupling of CZ neuron firing to cortical slow oscillations (Fig.3*G*–*J*). The similarities in the firing of CZ neurons in dopamine-intact and lesioned rats were evident in linear phase histograms (Fig.3*G*,*H*) and circular plots (Fig.3*I*,*J*). Moreover, the mean angles of firing of phase-locked CZ neurons in control and lesioned rats were not shifted (*p* = 0.560, common median test). When comparing vector lengths (Fig.3*K*), which indicate how spiking activity of single neurons is concentrated around a given preferred phase, it was evident that BZ neurons in dopamine-intact rats had more consistent phase-locked firing than neurons in the other three groups, which were all similar (*p* = 6.30×10^-8^, χ^2^ = 36.35, Kruskal-Wallis ANOVA; *p* = 0.004 for BZ intact vs BZ lesioned, *p* = 4.18×10^-8^ for BZ intact vs CZ intact, *p* = 0.008 for BZ intact vs CZ lesioned, *p* = 1.00 for BZ lesioned vs CZ intact, *p* = 1.00 for BZ lesioned vs CZ lesioned, *p* = 1.00 for CZ intact vs CZ lesioned, *post hoc* Dunn’s tests).

**Figure 3.**
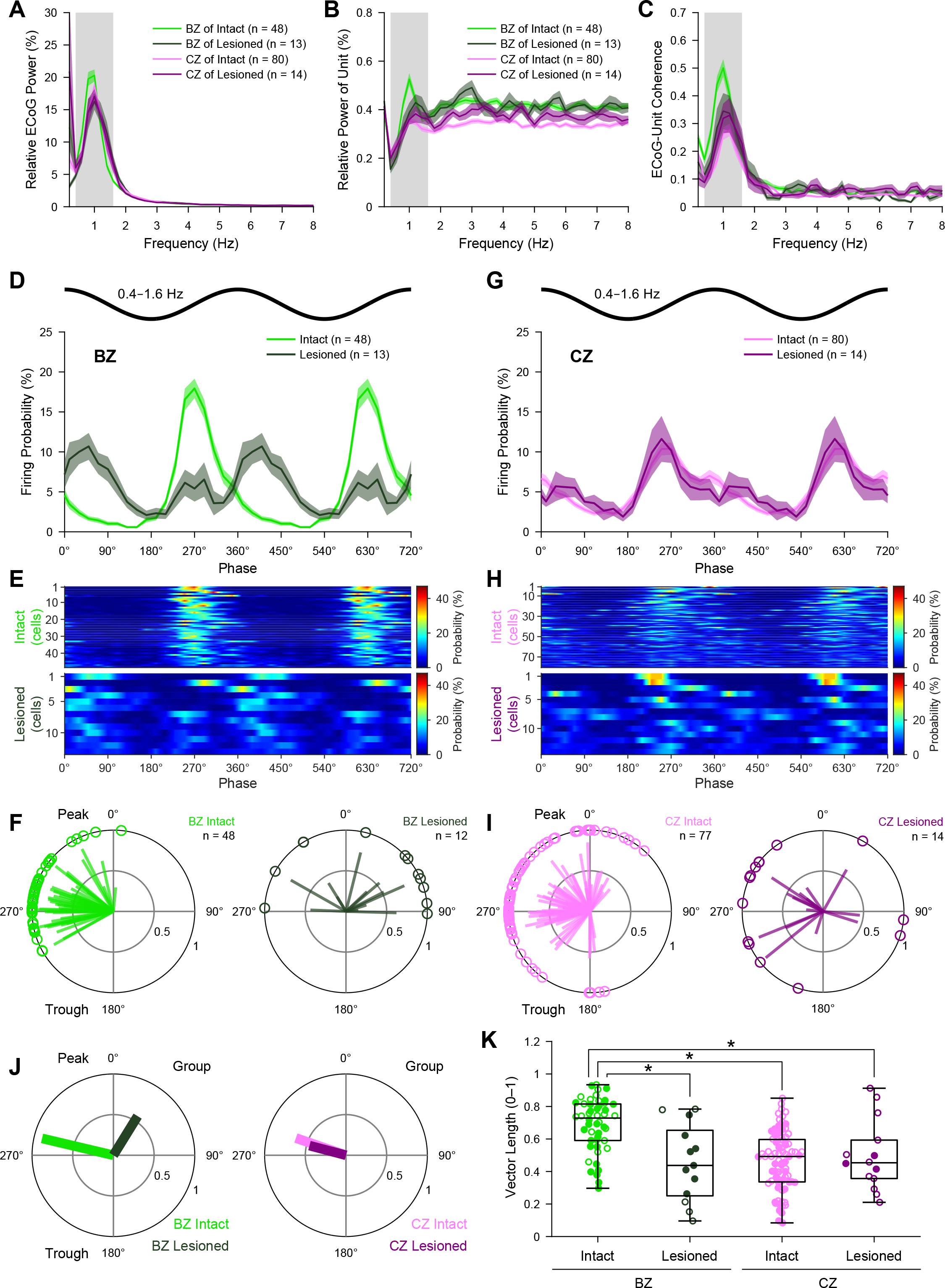
Spike timings of BZ and CZ neurons in relation to cortical slow oscillations. ***A***, Mean power spectra of ECoGs simultaneously recorded with all BZ neurons and all CZ neurons in dopamine-intact and 6-OHDA-lesioned rats. Power is relative to that at 0.4–100 Hz. Gray shading denotes frequency band of slow oscillations analyzed (0.4–1.6 Hz). ***B***, Mean power spectra of the spike discharges of all BZ neurons and CZ neurons. Note the peaks in power at frequencies similar to those of the cortical slow oscillations (***A***). ***C***, Mean coherence spectra between ECoGs and all BZ neurons and CZ neurons (color coding and sample sizes as in ***B***). Bin size of spectra in ***A–C*** is 0.2 Hz. ***D***, Linear phase histograms of the spike discharges of all BZ neurons with respect to cortical slow oscillations (bin size = 20°). Note the altered temporal coupling after dopamine depletion. For clarity, two cortical slow oscillation cycles are shown. ***E***, Heat map representation of linear phase histograms of individual BZ neurons (bin size = 20°; neurons sorted by vector length, with longest at top). ***F***, Circular plots of phase-locked firing of BZ neurons in dopamine-intact and lesioned rats. In this and subsequent circular plots of individual neurons, only those neurons that qualified for analyses (i.e. ý40 spikes recorded) and were significantly phase-locked (*p* < 0.05, Rayleigh’s Uniformity Test) are shown. Vectors of preferred firing of individual neurons are shown as lines radiating from the center. Greater vector lengths indicate lower variance in the distribution around the mean phase angle. Each circle on the plot perimeter represents the preferred phase (i.e. mean phase of all the spikes) of an individual neuron. ***G–I***, As in ***D–F***, but for CZ neurons. ***J***, Mean vectors of the preferred firing phases of thalamic neurons in each group. Note that the angular shift in the vector of BZ neurons in lesioned rats. ***K***, Vector lengths for all the spikes of all BZ and CZ neurons in dopamine-intact and lesioned rats. Data from individual identified and extrapolated neurons are shown as filled and open circles, respectively. \**p* <0.05 (exact *p* values are in Results text), Dunn’s tests following Kruskal–Wallis ANOVA. Data in ***A–D*** and ***G*** are mean ± SEM. *n*, the number of neurons/ECoGs analyzed. All individual neurons recorded with glass electrodes.

Together, these data show that the phase-locked firing of BZ neurons, but not CZ neurons, is impaired and inappropriately timed with respect to cortical slow oscillations after dopamine depletion. As such, these results collectively define one manifestation of an input zone-selective dysrhythmia in motor thalamus.

### The firing of SNr neurons is aberrantly phase-locked to cortical slow oscillations after dopamine depletion

GABAergic SNr neurons innervate the BZ (Kuramoto et al., 2011), and their firing during SWA is altered by dopamine depletion (Belluscio et al., 2003; Tseng et al., 2005; Walters et al., 2007), together raising the possibility that the dysrhythmic firing of BZ neurons (Fig.3) is mediated by SNr neurons. However, it is unclear whether the timing of BZ and SNr activity would support such a relationship. To address this, we used silicon probes to record single-unit activity in the SNr of dopamine-intact and 6-OHDA-lesioned rats during SWA (Fig.4). Although the power of ECoG slow oscillations simultaneously recorded with SNr neurons was slightly lower in lesioned rats (*p* = 0.016, MWUT; Fig.4*A*), the power of slow oscillations in SNr unit activities was greatly elevated in lesioned rats (*p* < 10^-9^, MWUT; Fig.4*B*). Moreover, qualitative inspection of SNr unit activities (Fig.4*D*), and analyses of phase histograms (Fig.4*E*,*F*), suggested that the temporal coupling of SNr neuron firing to cortical slow oscillations was markedly stronger in lesioned rats. Consistent with this, the firing of 88.4% of SNr neurons (38 of 43) in lesioned rats was significantly phase-locked (*p* < 0.05, Rayleigh’s Uniformity Test) to cortical slow oscillations, whereas only 48.7% of SNr neurons (18 of 37) were significantly phase-locked in dopamine-intact rats (Fig.4*J*,*K*). Moreover, the vector lengths of individual SNr neurons were longer in lesioned rats (*p* = 5.06×10^-5^, MWUT; Fig.4*L*), demonstrating that SNr neuron firing is more consistently phase-locked after dopamine depletion. These alterations in temporal coupling were not associated with changes in SNr neuron firing rates during SWA (18.81 ± 1.72 and 21.12 ± 0.21 spike/s in lesioned and dopamine-intact rats, respectively; *p* = 0.091, MWUT).

**Figure 4.**
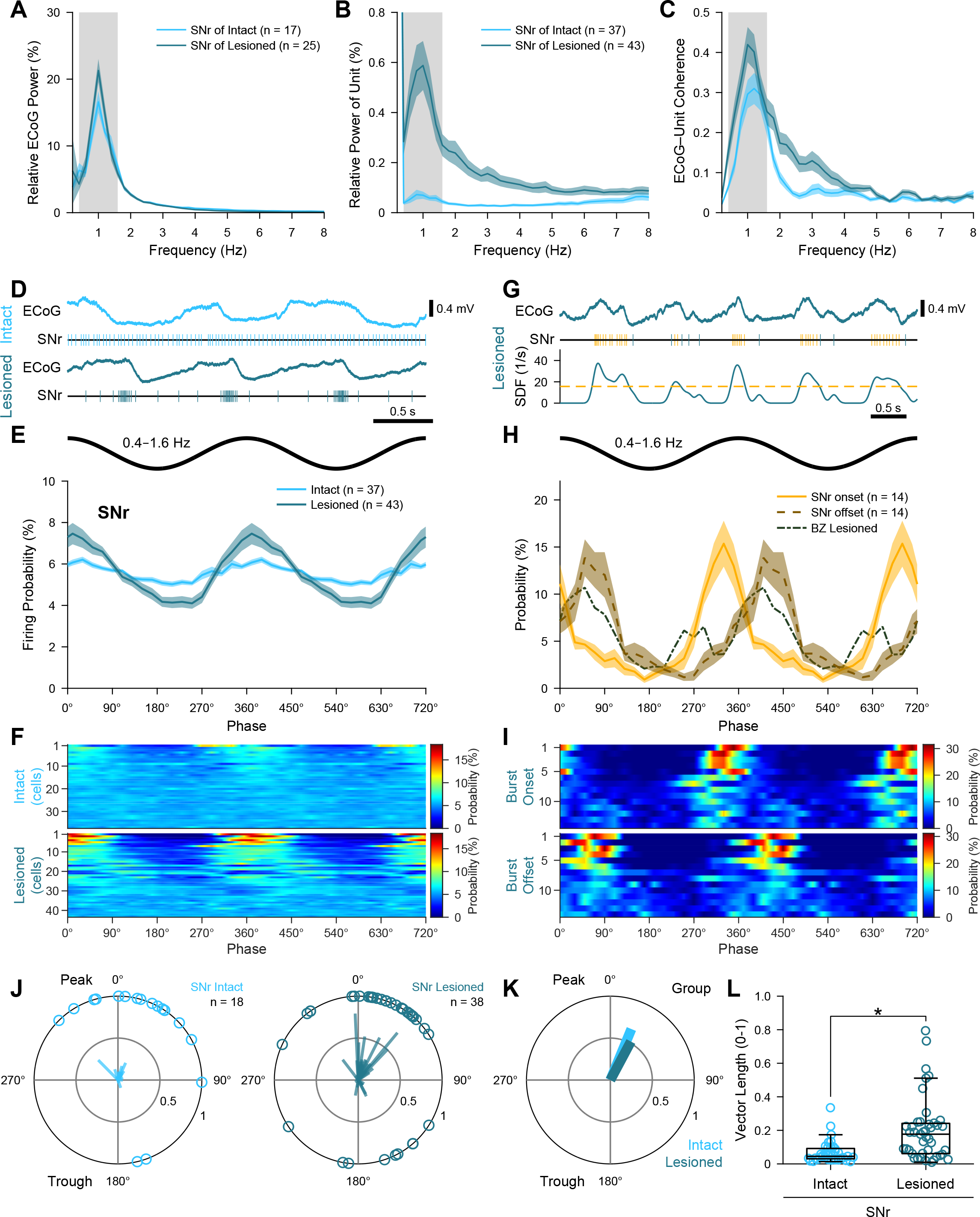
Spike timings of SNr neurons in relation to cortical slow oscillations. ***A***, Mean power spectra of ECoGs simultaneously recorded with SNr neurons in dopamine-intact and 6-OHDA-lesioned rats. Gray shading denotes frequency band of slow oscillations analyzed (0.4–1.6 Hz). ***B***, Mean power spectra of the spike discharges of all SNr neurons. Note the peaks in power at frequencies similar to those of the cortical slow oscillations (***A***). ***C***, Mean coherence spectra between ECoGs and all SNr neurons (color coding and sample sizes as in ***B***). Bin size of spectra in ***A–C*** is 0.2 Hz. ***D***, Example spike trains of single SNr neurons, shown together with simultaneously-recorded ECoGs for context, in an intact rat and a lesioned rat. ***E***, Linear phase histograms of the spike discharges of all SNr neurons with respect to cortical slow oscillations (bin size = 20°). Note the increased temporal coupling after dopamine depletion. ***F***, Heat map representation of linear phase histograms of individual SNr neurons (bin size = 20°; neurons sorted by vector length, with longest at top). ***G***, Detection of bursts of spikes fired by SNr neurons in lesioned rats. The spike train of a single SNr neuron (middle) and its spike density function (SDF, bottom) are shown together with the simultaneously-recorded ECoG (top). The yellow dashed line in the SDF plot indicates the threshold (see Materials and Methods) for detecting the onsets and offsets of bursts. Yellow and blue ticks in the spike train indicate spikes fired within and outside of bursts, respectively. ***H***, Linear phase histograms of the onsets (solid line in yellow) and offsets (dashed line in brown) of bursts fired by SNr neurons (*n* = 14; bin size = 20°) in lesioned rats, in comparison with the firing of BZ neurons in lesioned rats (dot and dashed line in gray). ***I***, Heat map representation of linear phase histograms of burst onsets and burst offsets of the 14 SNr neurons (bin size = 20°; neurons sorted by vector length). ***J***, Circular plots of phase-locked firing of individual SNr neurons in dopamine-intact and lesioned rats. ***K***, Mean vectors of the preferred phases of SNr neurons in each group. ***L***, Vector lengths for all the spikes of all SNr neurons in dopamine-intact and lesioned rats. \**p* = 5.06×10^-5^, Mann–Whitney U test. Data in ***A–C***, ***E*** and ***H*** are mean ± SEM. *n*, the number of neurons/ECoGs analyzed. All neurons are single units recorded with silicon probes.

On average, SNr neurons in both dopamine-intact and lesioned rats tended to fire just after the peak of cortical slow oscillations (Fig.4*E*,*J*,*K*); the mean angles of firing were 22.4 ± 11.2° and 28.1 ± 9.1°, respectively, and they were similar (*p* = 0.501, common median test; Fig.4*K*). Given that BZ neurons in lesioned rats fired at a mean angle of 31.7 ± 15.4° (Fig.3*J*), the average peak activity of SNr neurons would appear poorly timed to influence (inhibit) BZ neurons and shift their preferred phase of firing. However, we noted that some SNr neurons fired in a phasic ‘bursting’ manner, rhythmically alternating between intense firing and near quiescence (Fig.4*F*), rather than quasi-sinusoidal fluctuations in firing. To investigate this, we first defined the bursts fired by a group of SNr neurons (*n* = 14), chosen for their high firing variability (CV ≥ 1.0), and analyzed the timing of their burst onsets and offsets with respect to cortical slow oscillations (Fig.4*G–I*). Comparisons of linear phase histograms revealed that SNr burst onsets preferentially occurred just before the peak of cortical slow oscillations, and overlapped with reductions in BZ neuron activity, whereas SNr burst offsets preferentially occurred during the descending phase of cortical slow oscillations, and overlapped with the primary peak of BZ neuron activity (Fig.4*H,I*). These correlations suggest that, upon dopamine depletion, the emergence of aberrantly phase-locked burst firing in SNr is a valid candidate for mediating the phase shift and dysrhythmia of BZ neurons.

### The firing of individual BZ neurons, but not CZ neurons, is often aberrantly phase-locked to exaggerated cortical beta oscillations after dopamine depletion

We next examined the extent to which the firing of individual neurons (identified and extrapolated) in motor thalamus is altered with respect to the cortical beta-frequency (15–30 Hz) oscillations present during cortical activation (Fig.5; also see Fig.1*C,D*,*G,H*). In line with previous reports (Mallet et al., 2008a, 2008b; Sharott et al., 2017), the ECoGs simultaneously recorded with thalamic neurons in 6-OHDA-lesioned rats showed significantly exaggerated beta oscillations as compared to those recorded in dopamine-intact rats (*p* = 6.08×10^-6^ for ECoGs with BZ neurons, *p* = 3.67×10^-4^ for CZ neurons, MWUT; Fig.5*A*; also see Fig.1*D*,*H*). The power spectra of spikes fired by BZ neurons, but not CZ neurons, displayed significantly enhanced and prominent beta oscillations after dopamine depletion (*p* = 4.55×10^-5^ for BZ neurons in lesioned vs dopamine-intact rats, *p* = 0.291 for CZ neurons, MWUT; Fig.5*B*). Beta-band coherence between ECoGs and BZ neurons was significantly augmented after dopamine depletion (*p* = 1.03×10^-6^, MWUT; Fig.5*C*), but this was not the case for CZ neurons (*p* = 0.060, MWUT; Fig.5*C*). Phase histograms further suggested that the temporal coupling of BZ neuron firing to cortical beta oscillations was markedly stronger in lesioned rats (Fig.5*D,E*). Consistent with this, the firing of all BZ neurons (11 of 11) in lesioned rats was significantly phase-locked (*p* < 0.05, Rayleigh’s Uniformity Test) to cortical beta oscillations, whereas only 6.3% of BZ neurons (2 of 32) were significantly phase-locked in dopamine-intact rats. Changes in the temporal coupling of CZ neurons to cortical beta oscillations were less marked (Fig.5*F*,*G*), with minor proportions of CZ neurons exhibiting significantly phase-locked firing (3.3% [1 of 30] and 37.5% [3 of 8] of CZ neurons in dopamine-intact and lesioned rats, respectively). The disparities between BZ and CZ were unlikely to have arisen from systematic differences in cortical activity; the power of ECoG beta oscillations recorded with BZ or CZ neurons in lesioned rats was similar (*p* = 0.075, MWUT). In lesioned rats, BZ neurons tended to discharge during the descending phase of cortical beta oscillations (Fig.5*D*), with a mean angle of firing of 138.8 ± 12.3° for the group (Fig.5*H*). When comparing the vector lengths of individual neurons (Fig.5*I*), BZ neurons in lesioned rats had more consistent phase-locked firing than neurons in the other three groups, which were all similar (*p* = 2.57×10^-6^, χ^2^ = 28.70 Kruskal-Wallis ANOVA; *p* = 1.77×10^-6^ for BZ intact vs BZ lesioned, *p* = 1.00 for BZ intact vs CZ intact, *p* = 0.977 for BZ intact vs CZ lesioned, *p* = 7.96×10^-6^ for BZ lesioned vs CZ intact, *p* = 0.007 for BZ lesioned vs CZ lesioned, *p* = 0.997 for CZ intact vs CZ lesioned, *post hoc* Dunn’s tests).

**Figure 5.**
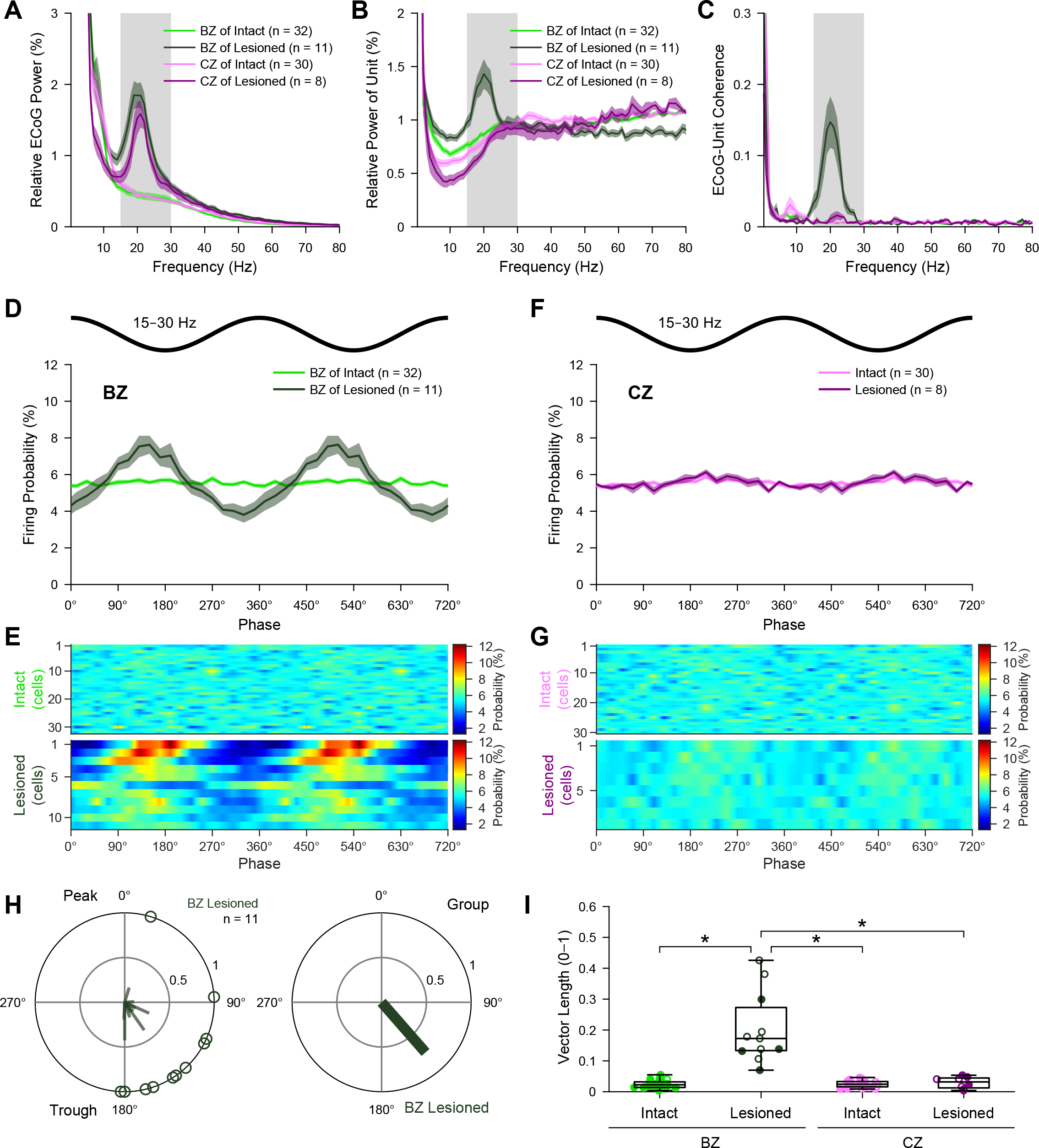
Spike timings of BZ and CZ neurons in relation to cortical beta oscillations. ***A***, Mean power spectra of ECoGs simultaneously recorded with all BZ neurons and all CZ neurons in dopamine-intact and 6-OHDA-lesioned rats. Power is relative to that at 1–100 Hz. Gray shading denotes frequency band of beta oscillations analyzed (15–30 Hz). ***B***, Mean power spectra of the spike discharges of BZ neurons and CZ neurons. Note the peak in power at beta frequencies for BZ neurons in lesioned rats; their peak frequency was similar to that of cortical beta oscillations (***A***). ***C***, Mean coherence spectra between ECoGs and all BZ neurons and CZ neurons (color coding and sample sizes as in ***B***). Note the peak in coherence at beta frequencies for BZ neurons in lesioned rats. Bin size of spectra in ***A–C*** is 1 Hz. ***D***, Linear phase histograms of the spike discharges of all BZ neurons with respect to cortical beta oscillations (bin size = 20°). Note the increased temporal coupling after dopamine depletion. For clarity, two cortical beta oscillation cycles are shown. ***E***, Heat map representation of linear phase histograms of individual BZ neurons (bin size = 20°; neurons sorted by vector length, with longest at top). ***F***, ***G***, As in ***D***, ***E***, but for CZ neurons. ***H***, *Left*, Circular plot of the phase-locked firing of individual BZ neurons in lesioned rats. *Right*, Mean vector of the preferred phases of BZ neurons in lesioned rats. ***I***, Vector lengths for all the spikes of all BZ and CZ neurons in dopamine-intact and lesioned rats. Data from individual identified and extrapolated neurons are shown as filled and open circles, respectively. \**p* <0.05 (exact *p* values are in Results text), Dunn’s tests following Kruskal–Wallis ANOVA. Data in ***A–D***, and ***F*** are mean ± SEM. *n*, the number of neurons/ECoGs analyzed. All individual neurons recorded with glass electrodes.

Taken together, these data show that individual BZ neurons, but not CZ neurons, tend to inappropriately engage with the exaggerated cortical beta oscillations that arise during activated brain states after dopamine depletion. As such, these results collectively reveal a second manifestation of an input zone-selective dysrhythmia in motor thalamus.

### Neuronal ensemble activity in the BZ, but not CZ, is aberrantly synchronized and phase-locked to exaggerated cortical beta oscillations after dopamine depletion

To gain insight into whether and how alterations in motor thalamus activity occurring after dopamine depletion extended to the collective outputs from larger ensembles of neurons, we analyzed silicon probe recordings of background-unit activity (BUA) that represents the spike firing of many neurons around the probe contacts (Moran et al., 2008; Moran and Bar-Gad, 2010). The BUA signals were extracted from wide-band recordings (Fig.6-1) and then used as continuous time series for spectral and circular statistical analyses. The use of DiI-coated linear probes with multiple recording contacts of known separation, together with *post hoc* anatomical verification of probe placement, allowed us to map the spatial profile of ensemble activity across discrete regions of BZ and CZ (Fig.6*A,A’*). Simultaneous recordings of BUA signals across the two thalamic input zones in lesioned rats showed that BZ ensemble activity, but not CZ ensemble activity, was often conspicuously modulated in time with ongoing cortical beta oscillations (Fig.6*B*). Accordingly, some BUA signals in BZ, but not CZ, also exhibited prominent beta oscillations (Fig.6*C*) and clear beta-band coherence with ECoGs (Fig.6*D*). Of note, the pronounced temporal coupling, power and coherence of BUA signals at beta frequencies was not always present across the whole BZ, such that focal ‘hot spots’ were instead evident (Fig.6*B*–*D*).

**Figure 6.**
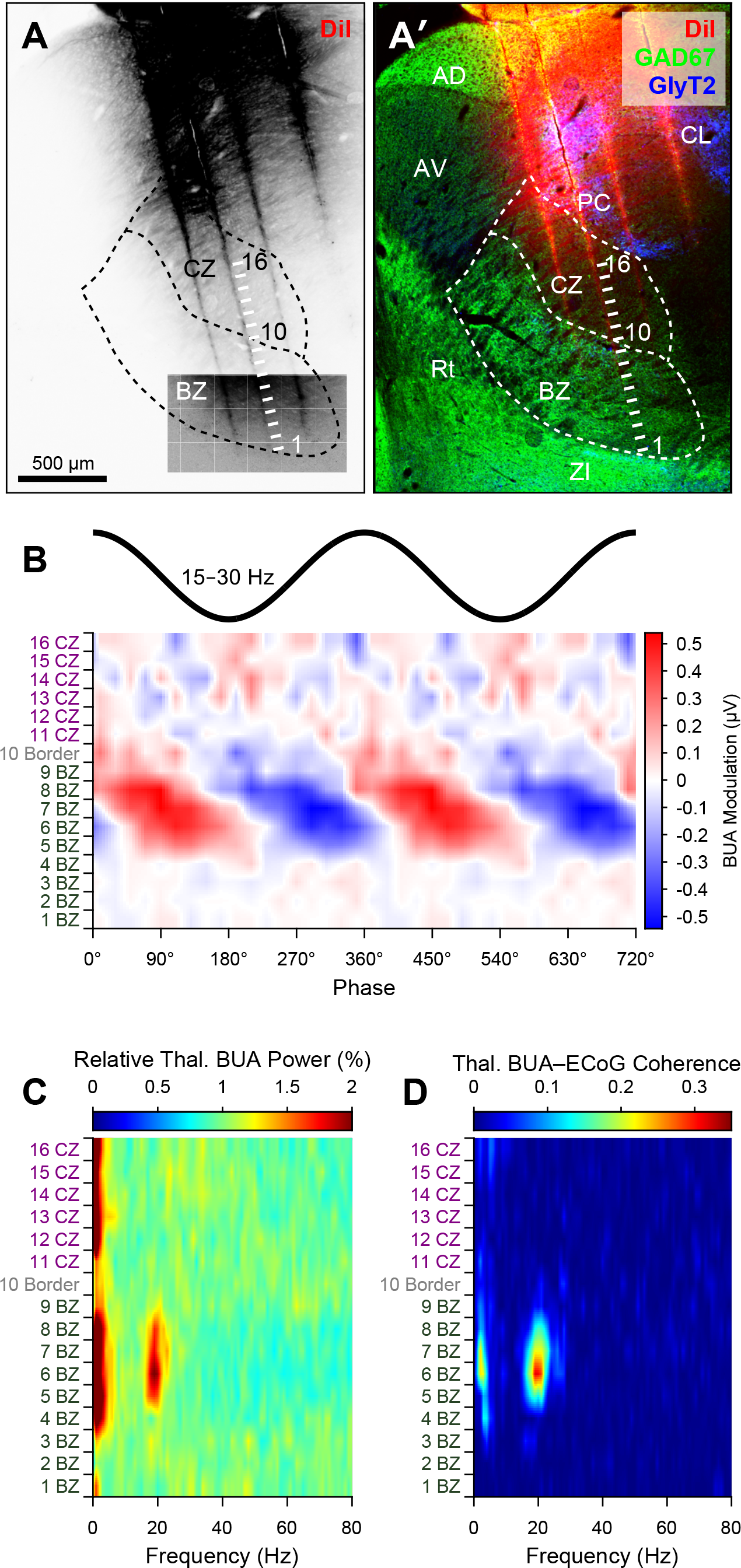
Example localization of background-unit activity signals in the motor thalamus, and their relationship with cortical beta oscillations. ***A***, Image of a parasagittal tissue section from a 6-OHDA-lesioned rat, with DiI fluorescence signal (in an inverted tone for clarity) marking four penetration tracks made at different times by a DiI-coated silicon probe during recordings *in vivo*. An overlaid image with enhanced contrast reveals the ends of three tracks. The estimated positions of the probe’s 16 recording contacts along one such track are denoted by short lines. ***A’***, The DiI signal (red) was localized with respect to the basal ganglia-recipient zone (BZ) and cerebellar-recipient zone (CZ) of the motor thalamus, as delineated by GAD67 (green) and GlyT2 (blue) immunofluorescence, on the same section as in ***A***. Probe contacts 1–9 were considered to have been within BZ, whereas contacts 11–16 were in CZ. ***B***, Background-unit activity (BUA) signals from the 16 recording contacts at the positions indicated in ***A/A’***, averaged according to the instantaneous phases of cortical beta oscillations (bin size = 20°) and plotted as a heat map. Note the large-amplitude phasic modulation of BUA signals at contacts 5–9 within the BZ. ***C***, Power spectra of BUA signals, from the same recordings as in ***B***, shown as a heat map. Power is relative to that at 1–100 Hz. ***D***, Spectra of coherence between the same BUA signals and the simultaneously-recorded ECoG. Note the focal nature of thalamic power and coherence at beta frequencies. Bin size of spectra in ***C*** and ***D*** is 1 Hz. In the parasagittal section shown in ***A/A’***, rostral is towards the left, and dorsal is towards the top. AD, anterodorsal thalamic nucleus; CL, central lateral thalamic nucleus; PC, paracentral thalamic nucleus; Rt, thalamic reticular nucleus; ZI, zona incerta. For abbreviations of other thalamic nuclei, see Fig1.

We extended the analysis of thalamic BUAs to quantitative comparisons across dopamine-intact and lesioned rats (*n* = 6 and 7 rats, respectively) (Fig.7). Among the power spectra of thalamic BUAs, only those recorded in the BZ of lesioned rats showed a conspicuous peak at beta frequencies (Fig.7*B*). Accordingly, beta-band power in BZ BUA signals was significantly increased after dopamine depletion (*p* = 2.87×10^-8^, MWUT). Similarly, the beta-band coherence between thalamic BUA signals and ECoGs was significantly increased for BZ, but not for CZ, after dopamine depletion (*p* = 9.99×10^-14^ for BZ, *p* = 0.444 for CZ, MWUT; Fig.7*C*). These disparities were unlikely to have arisen from systematic differences in cortical activity; the power of ECoG beta oscillations recorded with BZ or CZ BUA signals in lesioned rats was similar (*p* = 0.920, MWUT; Fig.7*A*). Beta-band coherence between pairs of BZ BUA signals was significantly increased after dopamine depletion (*p* = 2.08×10^-23^, MWUT; Fig.7*D*). However, this was not the case for pairs of CZ BUA signals (*p* = 0.901, MWUT; Fig.7*E*), nor for pairs made up of one BZ signal and one CZ signal (*p* = 0.834, MWUT; Fig.7*F*). As noted above, not all BZ BUA signals showed clear peaks in power or coherence at beta frequencies after dopamine depletion (Fig.6). Thus, to characterize the phase relationships between BZ BUA and cortical beta oscillations in lesioned rats, we first selected the BZ BUA signals with sample vector lengths longer than the 99.9th percentile of the sample vector lengths of the shuffled data (see Materials and Methods). When the amplitudes of these ‘significantly modulated’ BUA signals (*n* = 35) were plotted with respect to cortical beta oscillations, it was evident that BZ ensemble activity tended to increase during the descending phase (Fig.7*G*,*H*). On average, these BZ BUA signals showed a preferred mean angle of 123.4 ± 7.2° (Fig.7*I*), which is similar (*p* = 0.300, common median test) to the value we obtained for individual neurons in BZ (138.8 ± 12.3°; Fig.5*H*). The power spectrum of the significantly-modulated BZ BUA signals had a prominent peak in the beta band (Fig.7*J*), as did their coherence spectrum with ECoGs (Fig.7*K*).

**Figure 7.**
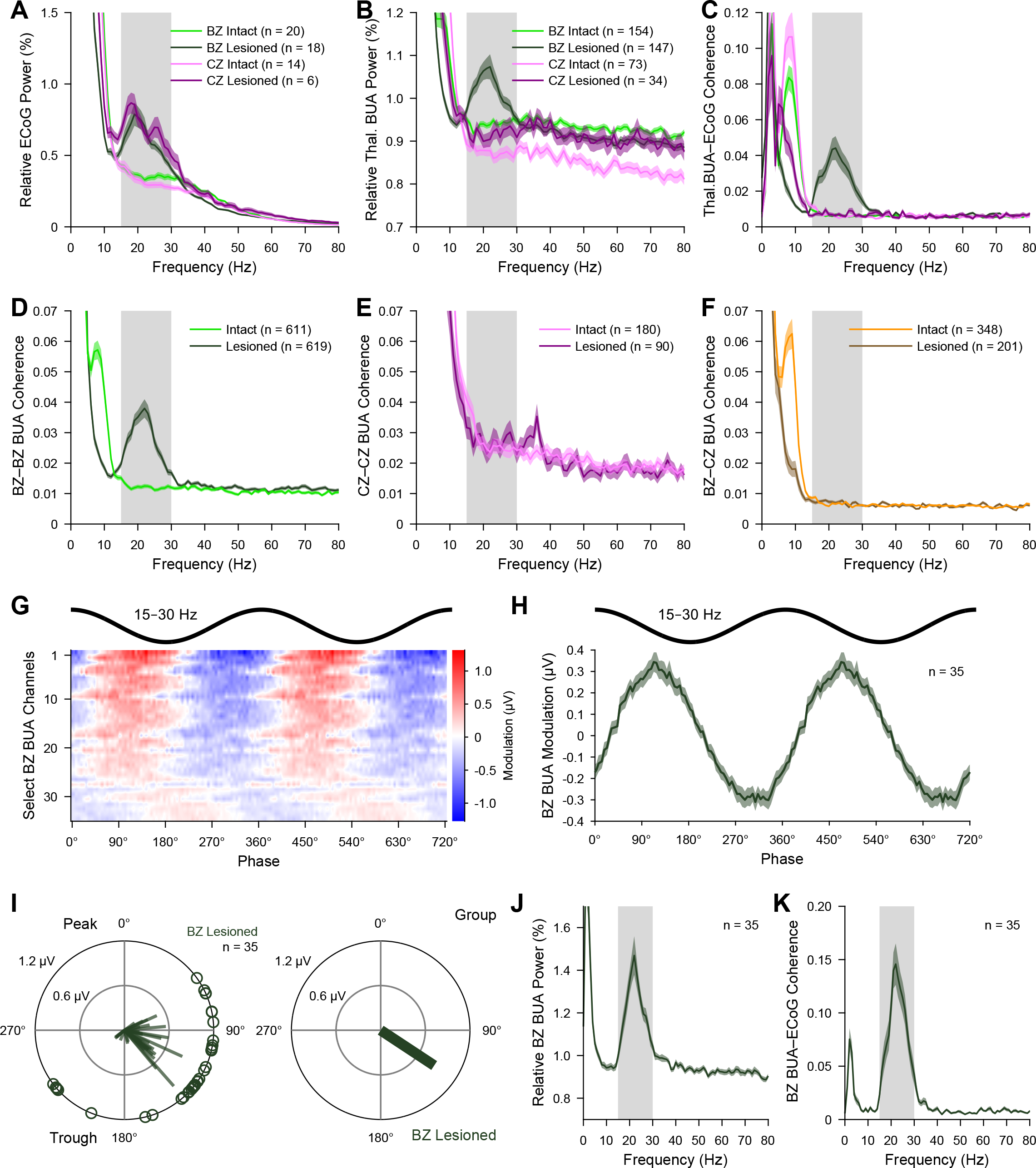
Background-unit activity signals in the motor thalamus during cortical activation in dopamine-intact and 6-OHDA-lesioned rats. ***A***, Mean power spectra of ECoGs simultaneously recorded with background-unit activity (BUA) in BZ and CZ in dopamine-intact and 6-OHDA-lesioned rats. Gray shading denotes frequency band of beta oscillations analyzed (15–30 Hz). ***B***, Mean power spectra of BUA signals recorded in motor thalamus. Note the peak in power at beta frequencies for the BZ BUA signals in lesioned rats. ***C***, Mean coherence spectra between the thalamic BUA signals and ECoGs (color coding and sample sizes as in ***B***). Note the peak in coherence at beta frequencies for BZ BUA signals in lesioned rats. ***D***, Coherence spectra of simultaneously-recorded pairs of BZ BUA signals. ***E***, As in ***D***, but for pairs of CZ BUA signals. ***F***, Coherence spectra between simultaneously-recorded pairs of BZ and CZ BUA signals. ***G***, Heat map representation of the phase-averaged waveforms of the significantly-modulated BZ BUA signals (***n*** = 35) in lesioned rats (bin size = 5°; BUA signals sorted by vector length). ***H***, Group average of waveforms shown in ***G***. Note that positive modulations of BZ BUA tended to occur during the descending phase of cortical beta oscillations. ***I***, *Left,* Circular plots of the individual significantly-modulated BZ BUA signals in lesioned rats. Vectors representing the phase preference of individual BUA signals are shown as lines radiating from the center. Greater vector lengths indicate greater modulation of BUA amplitude around the mean phase angle. Each circle on the plot perimeter represents the preferred phase of an individual BUA signal. *Right*, Mean vector of the preferred phases of all significantly-modulated BZ BUA signals. ***J***, Mean power spectrum of the significantly-modulated BZ BUA signals. ***K***, Mean coherence spectrum between significantly-modulated BZ BUA signals and the simultaneously-recorded ECoGs. Data in ***A–F***, ***H***, ***J*** and ***K*** are mean ± SEM. Bin size of power and coherence spectra is 1 Hz. ***n***, the number of individual ECoG recordings (***A***), the number of individual BUA signals recorded (***B***, ***C***), the number of pairs of BUA signals recorded (***D–F***), the number of the significantly-modulated BZ BUA signals (***G–K***). All BUA recorded with silicon probes.

In summary, these data collectively demonstrate that, after dopamine depletion, neuronal ensembles in the BZ, but not CZ, inappropriately synchronize their outputs at beta frequencies during cortical activation. Thus, input zone-selective dysrhythmia in motor thalamus also manifests at the level of population activity.

### Neuronal ensemble activity in the SNr is aberrantly synchronized and phase-locked to exaggerated cortical beta oscillations after dopamine depletion

To provide further context for the dysrhythmic firing of BZ neurons in relation to exaggerated beta oscillations, we next tested whether neuronal ensemble activity in SNr during cortical activation was similarly altered by dopamine depletion (Figs.8 and 9). Following the experimental approach used to investigate motor thalamus (Figs.6 and 7), the use of DiI-coated silicon probes, together with *post hoc* anatomical verification of probe placement, allowed us to map the spatial profile of activity across discrete regions of SNr (Fig.8*A,A’*). Simultaneous recordings of BUA signals at multiple sites within the SNr of lesioned rats showed that ensemble activity was often conspicuously modulated in time with ongoing cortical beta oscillations (Fig.8*B*). Some SNr BUA signals also exhibited prominent beta oscillations (Fig.8*C*) and clear beta-band coherence with ECoGs (Fig.8*D*). The pronounced temporal coupling, power and coherence of BUA signals at beta frequencies typically manifested as focal hot spots (Fig.8*B*–*D*).

**Figure 8.**
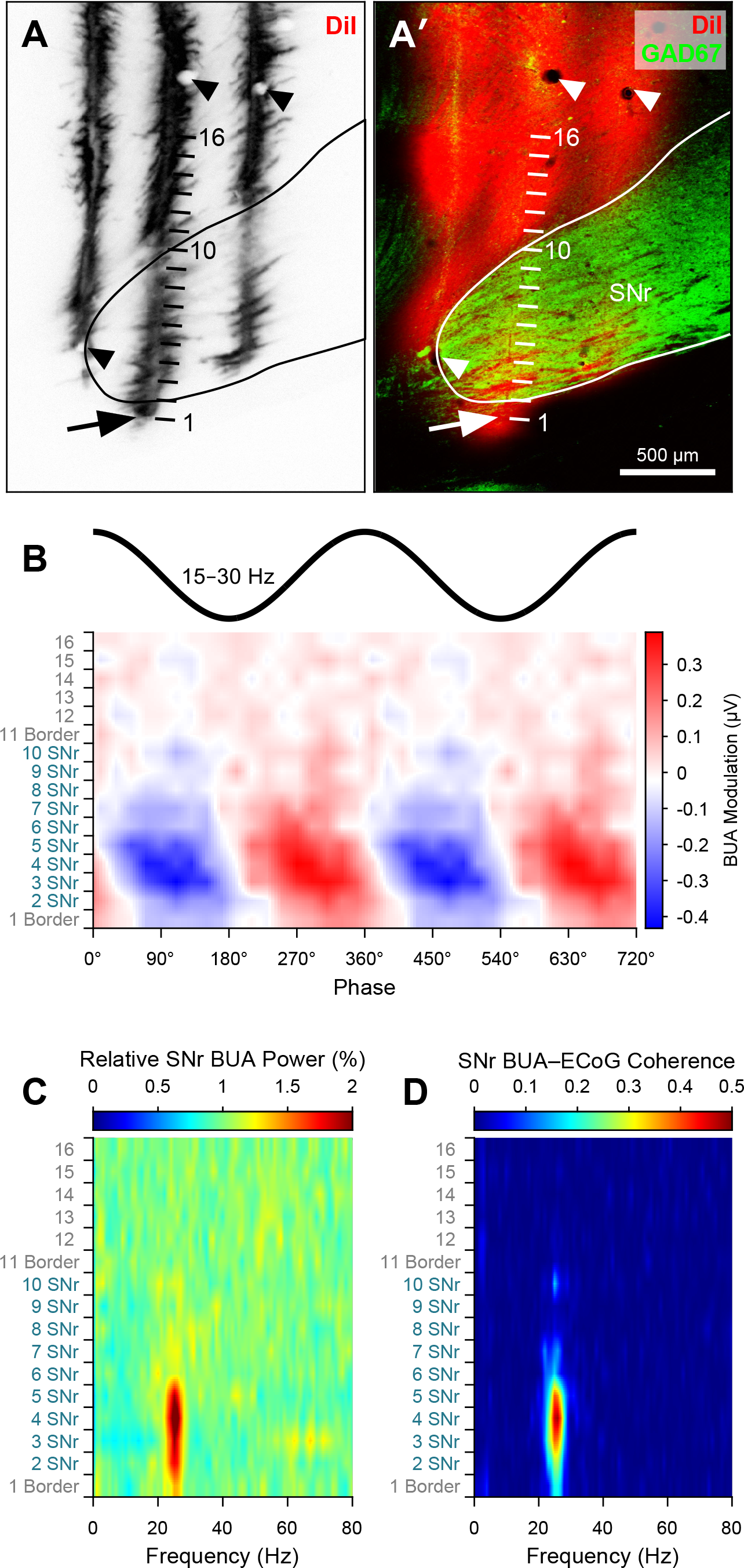
Example localization of background-unit activity signals in the SNr, and their relationship with cortical beta oscillations. ***A***, Image of a parasagittal tissue section from a 6-OHDA-lesioned rat, with DiI fluorescence signal (in an inverted tone for clarity) marking three penetration tracks made at different times by a DiI-coated silicon probe during recordings *in vivo*. The estimated positions of the probe’s 16 recording contacts along one such track are denoted by short lines. ***A’***, The DiI signal (red) was localized with respect to the SNr, as delineated by GAD67 immunofluorescence (green), on the same section as in ***A***. Probe contacts 2–10 were considered to have been within SNr. ***B***, Background-unit activity (BUA) signals from the 16 recording contacts at the positions indicated in ***A/A’***, averaged according to the instantaneous phases of cortical beta oscillations (bin size = 20°) and plotted as a heat map. Note the large-amplitude phasic modulation of BUA signals at contacts 2–6 within the SNr. ***C***, Power spectra of BUA signals, from the same recordings as in ***B***, shown as a heat map. Power is relative to that at 1–100 Hz. ***D***, Spectra of coherence between the same BUA signals and the simultaneously-recorded ECoG. Note the focal nature of SNr power and coherence at beta frequencies. Bin size of spectra in ***C*** and ***D*** is 1 Hz. In the parasagittal section shown in ***A***/***A’***, rostral is towards the left, and dorsal is towards the top.

Quantitative comparisons of SNr BUA signals recorded in dopamine-intact and lesioned rats (*n* = 5 and 5 rats, respectively) revealed several important differences (Fig.9). The power spectrum of SNr BUA signals in lesioned rats, but not dopamine-intact rats, showed a prominent peak at beta frequencies (Fig.9*B*). Accordingly, beta-band power in SNr BUA signals was significantly increased after dopamine depletion (*p* = 8.19×10^-9^, MWUT). The beta-band coherence between SNr BUA signals and ECoGs significantly increased after dopamine depletion (*p* = 5.41×10^-31^, MWUT; Fig.9*C*), as did the beta-band coherence between pairs of SNr BUA signals (*p* = 3.97×10^-50^, MWUT; Fig.9*D*). Because not all SNr BUA signals showed clear peaks in power or coherence at beta frequencies after dopamine depletion (Fig.8), we again selected a set of significantly-modulated SNr BUA signals (*n* = 92) for analyses of phase relationships between SNr ensemble activity and cortical beta oscillations in lesioned rats (Fig.9*G*–*I*). Ensemble activity in SNr tended to increase during the ascending phase of cortical beta oscillations (Fig.9*G*,*H*), with, on average, a preferred mean angle of 266.1 ± 2.3° (Fig.9*I*). The power spectrum of the significantly-modulated SNr BUA signals had a prominent peak in the beta band (Fig.9*J*), as did their coherence spectrum with ECoGs (Fig.9*K*). On comparing the rhythmic modulation (preferred phases) of SNr BUA signals (Fig.9*H*,*I*) and BZ BUA signals (Fig.7*H*,*I*), an ‘anti-phase’ relationship was apparent, such that peaks in SNr ensemble activity were approximately timed with troughs in BZ ensemble activity, and *vice versa*. We also isolated single units from the same probe recordings in SNr. Beta-band coherence between SNr single units and ECoGs was significantly augmented after dopamine depletion (*p* = 6.03×10^-6^, MWUT; Fig.9*E*). The firing of 52% of SNr single units (24 of 46) in lesioned rats was significantly phase-locked (*p* < 0.05, Rayleigh’s Uniformity Test) to cortical beta oscillations, whereas none of the SNr single units (0 of 23) in dopamine-intact rats were similarly phase-locked. On average, SNr single units in lesioned rats fired at a preferred mean angle of 221.9 ± 11.1° (Fig.9*F*). These alterations in temporal coupling were not associated with changes in SNr neuron firing rates during cortical activation (29.25 ± 2.06 and 28.08 ± 0.32 spike/s in lesioned and dopamine-intact rats, respectively; *p* = 0.736, MWUT).

**Figure 9.**
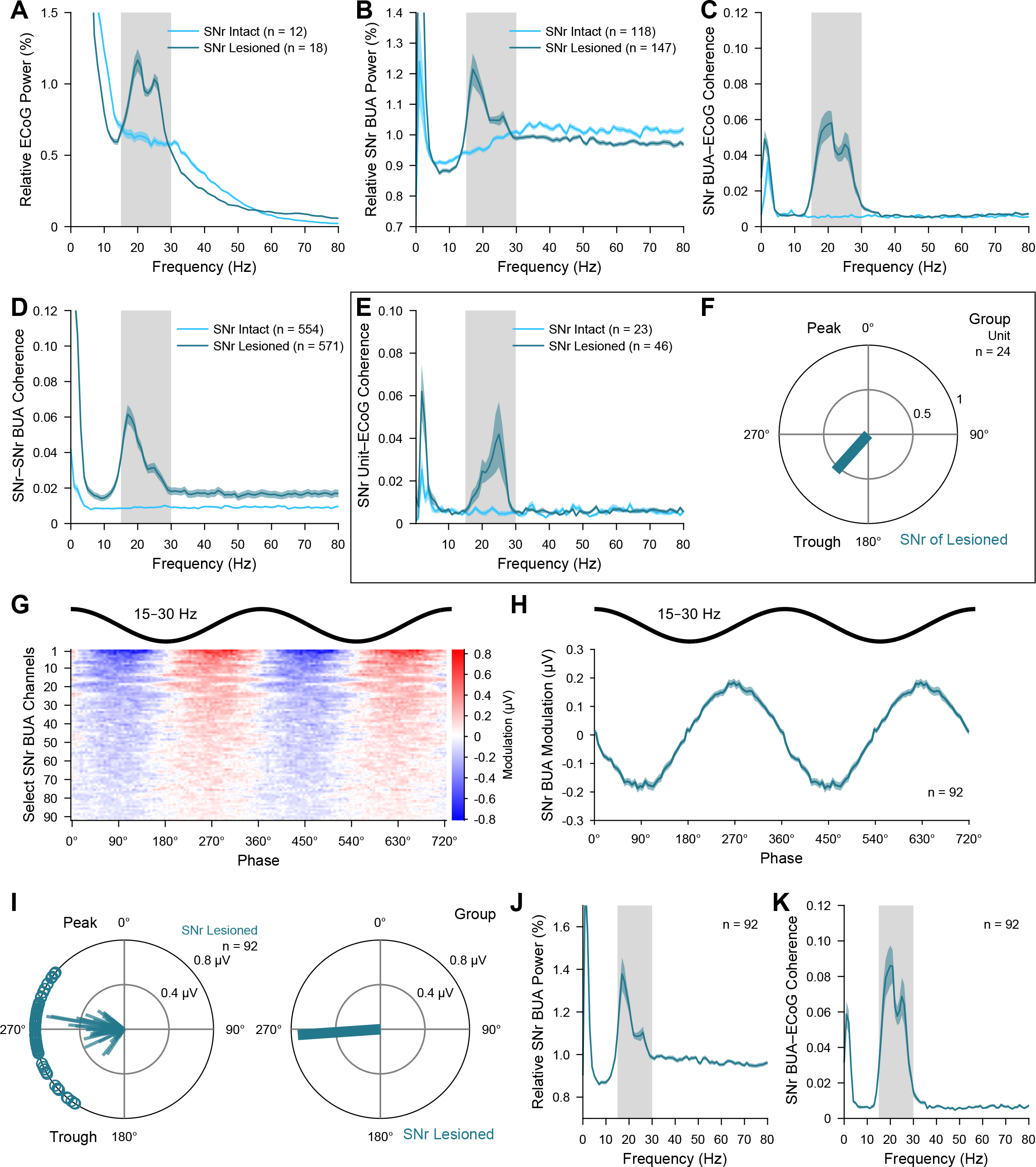
Background-unit activity signals in the SNr during cortical activation in dopamine-intact and 6-OHDA-lesioned rats. ***A***, Mean power spectra of ECoGs simultaneously recorded with background-unit activity (BUA) in the SNr in dopamine-intact and 6-OHDA-lesioned rats. Gray shading denotes frequency band of beta oscillations analyzed (15–30 Hz). ***B***, Mean power spectra of BUA signals recorded in SNr. Note the peak in power at beta frequencies for the SNr BUA signals in lesioned rats. ***C***, Mean coherence spectra between the SNr BUA signals and ECoGs (color coding and sample sizes as in ***B***). Note the peak in coherence at beta frequencies for SNr BUA signals in lesioned rats. ***D***, Coherence spectra of simultaneously-recorded pairs of SNr BUA signals. ***E***, Mean coherence spectra between SNr single units and ECoGs. ***F***, Mean vector of the preferred phases of SNr single units in lesioned rats. ***G***, Heat map representation of the phase-averaged waveforms of the significantly-modulated SNr BUA signals (*n* = 90) in lesioned rats (bin size = 5°; BUA signals sorted by vector length). ***H***, Group average of waveforms shown in ***G***. Note that positive modulations of SNr BUA tended to occur during the ascending phase of cortical beta oscillations. ***I***, *Left,* Circular plots of the individual significantly-modulated SNr BUA signals in lesioned rats. *Right*, Mean vector of the preferred phases of all significantly-modulated SNr BUA signals. ***J***, Mean power spectrum of the significantly-modulated SNr BUA signals. ***K***, Mean coherence spectrum between significantly-modulated SNr BUA signals and the simultaneously-recorded ECoGs. Data in ***A–E***, ***H***, ***J*** and ***K*** are mean ± SEM. Bin size of power and coherence spectra is 1 Hz. *n*, the number of individual ECoG recordings (***A***), the number of individual BUA signals recorded (***B***, ***C***), the number of pairs of BUA signals recorded (***D***), the number of single units (***E***), the number of significantly phase-locked single units (***F***), the number of the significantly-modulated SNr BUA signals (***G–K***). All BUA signals and single units were recorded with silicon probes.

In summary, these data demonstrate that, after dopamine depletion, neuronal ensembles in the SNr inappropriately synchronize their outputs at beta frequencies during cortical activation. They further suggest that aberrantly phase-locked firing of SNr neurons is a valid candidate for mediating the related dysrhythmia of BZ neurons.

### GABA infusions into motor thalamus reduce Parkinsonian beta oscillations in motor cortex

Excessively synchronized beta-frequency output from the BZ (Figs.5-7) might be causally important for the expression of abnormal beta oscillations in the BZ’s principal target, namely the motor cortex. We reasoned that, if this were the case, then suppressing BZ neuron activity would reduce the power of abnormal beta oscillations in motor cortex. To test this, we recorded ECoG beta oscillations during cortical activation in lesioned rats (*n* = 10) before, during and after infusions of a small volume (60 nl) of GABA solution (0.5 M) via a glass pipette inserted into the motor thalamus (Fig.10). We chose to microinfuse GABA (rather than other agonists of GABA receptors) because the inactivation effects of similarly small infusions of GABA typically abate within a few minutes (Kojima and Doupe, 2009); this in turn allowed us to perform repeated infusions (using a minimum interval of 10 mins) at one or more thalamic sites in a single animal (Fig.10*A*,*B*). Using this approach, we thus aimed to rapidly, but transiently, quash or otherwise perturb neuronal activity in a small volume of thalamic tissue. By marking the trajectories of the same glass micropipettes with synthetic fluorescent markers, we were able to accurately localize the GABA infusion sites to the BZ or CZ (Fig.10*C*). Our approach is to be contrasted with that used in a previous study (Brazhnik et al., 2016), where much larger volumes (∼10×) of the longer-lasting GABAA receptor agonist muscimol were injected via much larger cannulas (>10× in diameter) into the ventral medial (VM) nucleus and thereabouts, with the effects of drug injection being quantified hours after the event.

**Figure 10.**
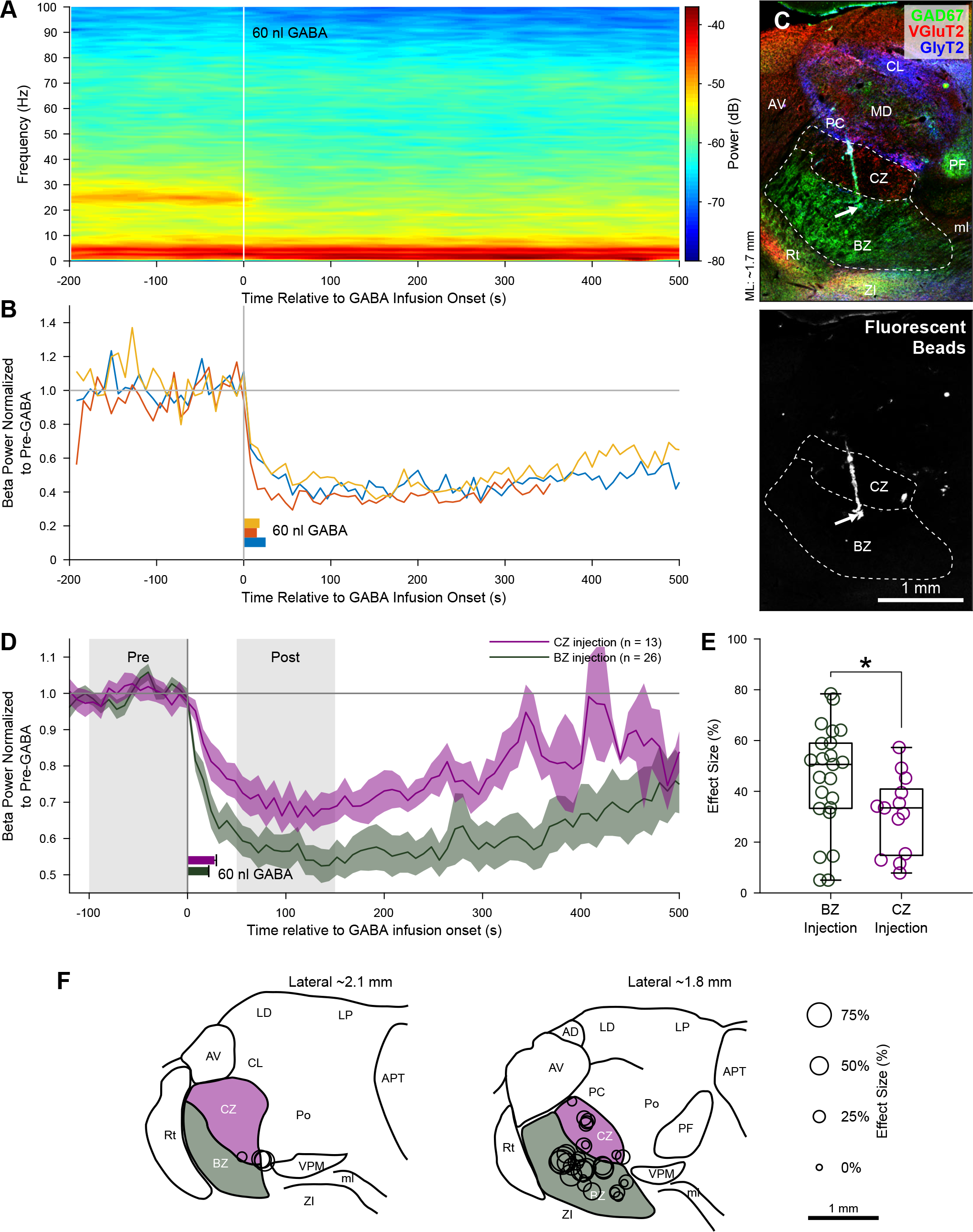
Effects of microinfusions of GABA into motor thalamus on the expression of cortical beta oscillations in 6-OHDA-lesioned rats. ***A***, Time-evolving power spectrogram of oscillations recorded over the motor cortex in a lesioned rat before, during and after a single infusion of a GABA solution (60 nl, 0.5 M) into the BZ of the motor thalamus. Note the clear reduction in the power of the abnormal beta oscillations (centre frequency of ∼24 Hz) a few seconds after the onset of the GABA infusion (at time = 0 s). ***B***, Power of cortical oscillations in the beta-frequency band (15–30 Hz; normalized to beta power in the 100 s immediately preceding GABA infusion) before, during and after three separate infusions of GABA at the same site in BZ (yellow, red and blue traces). The blue trace corresponds to the data shown in (***A***). The interval between each infusion was >10 min. The horizontal bars (yellow, red, blue) indicate the corresponding duration of each manually-controlled GABA infusion. Note the highly-reproducible time course and magnitude of the reduction in cortical beta power upon GABA infusion. ***C***, The GABA infusion site accessed in (***A***) and (***B***) was localized to the BZ of motor thalamus by *post hoc* anatomical analyses; the fluorescent beads marking the trajectory of the infusion pipette traversed the border between CZ and BZ (dashed line), and then terminated in the BZ (arrows). ***D***, Mean power of cortical beta oscillations before, during and after all infusions of GABA at BZ sites (green trace) or at CZ sites (purple trace) in the motor thalamus. Green and purple shaded areas show SEMs. Horizontal bars indicate the mean durations (+ SEMs) of the manually-controlled GABA infusions into BZ (green) or CZ (purple). The nadirs in cortical beta power occurred around 100 s after the onset of GABA infusion. Note that the cortical beta power at 50–150 s after BZ infusions was lower than that after CZ infusions. ***E***, Effect sizes of all GABA infusions at BZ sites (green circles) or CZ sites (purple circles). Effect size was defined as the percentage reduction in cortical beta power at 50–150 s after GABA infusion (“Post” gray shading in ***D***) as compared to power during the 100 s immediately before infusion (“Pre” gray shading in ***D***). Note that, on average, the effect sizes of BZ infusions were larger than those of CZ infusions (\**p* = 0.0297, Mann–Whitney *U* test). ***F***, Effect size of each GABA infusion according to its location in BZ and CZ. In the parasagittal sections shown in (***C***) and (***F***), rostral is towards the left, and dorsal is towards the top. APT, anterior pretectal nucleus; LD, lateral dorsal nucleus, lateral posterior nucleus, ml, medial lemniscus; PF, parafascicular nucleus. For abbreviations of other structures in and around the thalamus, see Figs.1 and 6.

Brief microinfusions (24.4 ± 1.0 s) of GABA at sites *post hoc* localized to BZ caused rapid (within ∼25 s of infusion onset) and reproducible decreases in the power of cortical beta oscillations (Fig.10*A*–*C*). On average, the nadir in cortical beta power occurred around 100 s from the onset of GABA infusions at BZ sites (*n* = 32), after which beta power steadily increased (Fig.10*D*). Effect size, defined as the percentage reduction in cortical beta power at 50–150 s after GABA infusion as compared to power during the 100 s immediately before infusion, was on average ∼50% for infusions at BZ sites (Fig.10*D*–*F*). That said, effect sizes across different BZ infusions sites and all animals were highly variable (Fig.10*D–F*), which might relate to the focal hot spot expression of beta oscillations in the BZ (see Fig.6). Infusion of GABA at CZ sites (*n* = 23) also led to reductions in cortical beta power, with an average time course of effect that was akin to that after infusions at BZ sites (Fig.10*D*). However, compared to infusions at BZ sites, the effect sizes of infusions at CZ sites were on average significantly smaller (*p* = 0.0297, MWUT; Fig.10*E*). As a further distinction, effects sizes of >60 % were limited to GABA infusions made at BZ sites (Fig.10*E,F*).

Taken together, these pharmacological perturbation analyses suggest that ongoing activity in the motor thalamus is permissive for exaggerated beta oscillations in motor cortex, with output from the BZ being of special importance for the maintenance of these pathological cortical rhythms.

## Discussion

Here, we elucidate how chronic dopamine depletion alters the temporal dynamics of electrical activity within the two major input zones of motor thalamus *in vivo*. Our data demonstrate that distinct, brain state-dependent manifestations of dysrhythmia selectively emerge within the basal ganglia-recipient zone in Parkinsonism.

### Neuronal firing rates in Parkinsonism

The direct/indirect pathways model predicts that motor thalamus neurons are hypoactive in Parkinsonism (DeLong, 1990; Smith et al., 1998; Galvan et al., 2015). Our recordings of individual neurons accurately localized to the BZ and CZ do not support this prediction. Previous work in anesthetized rats shows that dopamine depletion only alters the firing rates of some BG neurons in the expected manner during certain brain states (Belluscio et al., 2003; Abdi et al., 2015; Sharott et al., 2017). We determined that, irrespective of two extreme brain states, the spontaneous firing rates of BZ neurons were not abnormally decreased. Our results agree with electrophysiological studies of BZ neurons in awake dopamine-intact and Parkinsonian monkeys (Pessiglione et al., 2005; Kammermeier et al., 2016), suggesting they generalize beyond anesthesia and across species. *Ex vivo* BZ neurons exhibit augmented ‘rebound’ LTS bursting after dopamine depletion (Bichler et al., 2021). Our data show this does not manifest as altered propensities to fire spikes in LTS bursts *in vivo*. That BZ neurons were not hypoactive tallies with our observation that SNr neurons were not hyperactive after dopamine depletion; the latter is a recurring finding in awake and anesthetized rodents (Tseng et al., 2005; Walters et al., 2007; Lobb and Jaeger, 2015; Willard et al., 2019). We observed, however, that CZ neurons had increased firing rates during cortical activation, adding to evidence of altered CZ activity in Parkinsonism (Galvan et al., 2015; Wichmann, 2019). This CZ hyperactivity might reflect augmented output from (presumably glutamatergic) neurons in cerebellar nuclei after dopamine depletion (Menardy et al., 2019). Our recordings in BZ do not uphold canonical firing rate-based models of basal ganglia–thalamocortical dysfunction in Parkinsonism, but the predicted alterations might still emerge during specific motor behaviors. Nevertheless, our data support and extend the concept that profoundly dysrhythmic activity can arise without firing rate changes in the Parkinsonian BZ and SNr.

### Dysrhythmia in the Parkinsonian motor thalamus during slow-wave activity

Previous studies of BG neurons show that dopamine depletion can alter the patterning of their discharges during, and with respect to, the stereotyped cortical slow oscillation, thereby providing valuable insights into the potential contributions of different sets of inputs to their activity (Magill et al., 2001; Belluscio et al., 2003; Tseng et al., 2005; Mallet et al., 2006; Walters et al., 2007; Zold et al., 2007, 2012; Abdi et al., 2015; Sharott et al., 2017). An extreme example of altered activity patterning occurs in the external globus pallidus; the firing of prototypic neurons in dopamine-intact animals tends to increase slightly around the peaks of cortical slow oscillations, whereas prototypic neuron firing in 6-OHDA-lesioned animals is strongly timed with slow oscillation troughs (Mallet et al., 2008a, 2012; Abdi et al., 2015). This aberrant ‘anti-phase’ oscillatory firing of prototypic neurons in Parkinsonism is likely the result of receiving hypersynchronous rhythmic GABAergic inputs from striatal neurons (Zold et al., 2012; Nevado-Holgado et al., 2014; Sharott et al., 2017; Kovaleski et al., 2020). It follows that hypersynchronous rhythmic GABAergic outputs from BG should aberrantly entrain thalamic neurons, such that their preferred phase of firing is disturbed (Tseng, 2009). One previous study has addressed this prediction, revealing that neurons in the thalamic parafascicular nucleus, also targeted by GABAergic BG outputs, do not exhibit the expected anti-phase firing (Parr-Brownlie et al., 2009). In stark contrast, we demonstrate here that this prediction holds true for BZ neurons, with dopamine depletion resulting in weaker phase-locking and a ∼100° shift in the mean angles of their firing with respect to cortical slow oscillations. Notably, the phase-locked firing of CZ neurons was not inappropriately timed to slow oscillations, reinforcing that SWA-related dysrhythmia in the Parkinsonian motor thalamus is selective for input zone. In line with other studies (Belluscio et al., 2003; Tseng et al., 2005; Walters et al., 2007; Lobb and Jaeger, 2015), the temporal coupling of SNr neuron firing to cortical slow oscillations was markedly stronger in lesioned rats, such that some SNr neurons fired in a phasic bursting manner. We determined that SNr burst onsets timed with cyclical reductions in BZ neuron firing, whereas SNr burst offsets timed with the primary peak of BZ neuron activity. As such, the aberrant phase-locked burst firing of SNr neurons appears well suited to mediate the phase shift and dysrhythmia of BZ neurons after dopamine depletion; in future studies, it would be important to progress from correlation to causation. Ascribing this role to SNr does not preclude the possibility that other inputs to BZ, such as those from cortex, play important roles in driving the dysrhythmic activity of BZ neurons. In turn, the dysrhythmic firing of BZ neurons would be broadcast to wide areas of frontal cortex (Herkenham, 1979; Kuramoto et al., 2009, 2015), wherein it might negatively impact on activity dynamics. Numerous studies in people with PD have demonstrated alterations in cortical SWA and slow-wave sleep, with further implications for symptoms and quality of life (Zahed et al., 2021).

### Dysrhythmia in the Parkinsonian motor thalamus during cortical activation

We observed that individual BZ neurons inappropriately engage with the exaggerated cortical beta oscillations that arise during activated brain states after dopamine depletion. This agrees with a study of the VM nucleus in awake 6-OHDA-lesioned rats (Brazhnik et al., 2016). It should be noted, however, that VM is difficult to objectively delineate with Nissl staining (as used in Brazhnik et al., 2016), and that VM is only a fraction of the BZ (Nakamura et al., 2014). Here, we provide important advances by accurately localizing recordings to the BZ and CZ (thus addressing selectivity for input zone), by defining the extent to which BZ neuronal ensembles are rhythmically synchronized, and by determining whether engagement in beta oscillations is accompanied by changes in BZ neuron firing rates. Our analyses of BUA signals show that, after dopamine depletion, focally-organized neuronal ensembles in the BZ inappropriately synchronize their outputs at beta frequencies. It was not possible to determine whether there was a phase shift in BZ neuron firing during cortical activation, as occurred during SWA, because so few neurons coupled their firing to the weak/transient cortical beta oscillations present in the dopamine-intact state. These experiments reveal a second manifestation of dysrhythmia in the Parkinsonian motor thalamus, and we reiterate that exaggerated beta oscillations arise in BZ without firing rate changes. Expression of pathological beta-frequency activities is exquisitely selective for input zone, such that CZ neurons are not similarly dysrhythmic. Studies of idiopathic PD support the notion that exaggerated beta-band synchronization of BG neuronal activity underpins bradykinesia/rigidity (Kühn et al., 2006, 2009; Ray et al., 2008; Sharott et al., 2014, 2018). We conclude that BZ neurons are primed to mediate the detrimental influences of abnormal beta-band activity on neuronal information processing and movement in PD.

We observed that the temporal coupling of cortical beta oscillations to the firing of individual SNr neurons was markedly stronger after dopamine depletion, as in awake rats (Brazhnik et al., 2012, 2014). Extending our analysis to BUA signals, we detail the novel observation that focal neuronal ensembles in the SNr inappropriately synchronize their outputs at beta frequencies. Importantly, there was an approximate anti-phase relationship between SNr and BZ ensemble activities at beta frequencies; on average, peaks in SNr ensemble activity preceded peaks in BZ ensemble activity by ∼217°. Given a range of beta oscillation periods of ∼33–66 ms, this phase difference would represent a time delay of ∼20–40 ms, which tallies with the time course of evoked nigrothalamic IPSPs and the subsequent pauses they cause in BZ neuron firing *ex vivo* (Edgerton and Jaeger, 2014). Together, our data suggest that aberrant inhibitory outputs arising from the hypersynchronized beta-band firing of SNr neurons is a valid candidate for orchestrating the related dysrhythmia of BZ neurons in Parkinsonism. Again, in future studies, it would be important to address causation.

Frontal cortical areas innervating motor thalamus exhibit exaggerated beta oscillations after dopamine depletion, as evidenced in our ECoG recordings here as well as those in awake rats (Sharott et al., 2005; Mallet et al., 2008b; Li et al., 2012; Brazhnik et al., 2016). It is thus possible that cortex also directly entrains BZ neurons to beta rhythms. However, it is unknown whether cortical neurons innervating BZ (and/or CZ) engage in abnormal beta oscillations. In terms of reciprocal influence, our GABA microinfusion experiments strongly suggest that ongoing activity in the BZ in particular bolsters exaggerated beta oscillations in motor cortex. This would fit well with BZ neurons being positioned, via extensive axon collateral networks (Kuramoto et al., 2009, 2015), to deliver beta-band inputs to many cortical neurons. We conclude that the dysrhythmic BZ is a critical node in the wider basal ganglia-thalamocortical loop circuit for the generation and/or maintenance of pathological cortical beta oscillations in PD.

## Acknowledgements

This work was supported by the Medical Research Council (MRC; Awards MC_UU_12020/5, MC_UU_12024/2, and MC_UU_00003/5 to P.J.M; Awards MC_UU_12024/1 and MC_UU_00003/6 to A.S.), Parkinson’s UK (Grant G-0806 to P.J.M.), and the Japan Society for the Promotion of Science Grants-in-Aid for Scientific Research (KAKENHI) (15H01663, 26430015). K.C.N. was supported in part by the Human Frontier Science Program (LT000396/2009-L). A.S. was supported in part by a Marie Curie European Re-integration Grant (SNAP-PD) awarded by the European Union. We thank: T. Kaneko for gifts of antibodies, and I. Bar-Gad for MATLAB code; N. Mallet for assistance in early stages of this work; R.W. Guillery, E. Kuramoto, F. Vinciati and H. Cagnan for insightful scientific discussions; and L. Conyers, J. Westcott, and B. Micklem for technical support.

---

**Figure 6-1 (Extended data).**
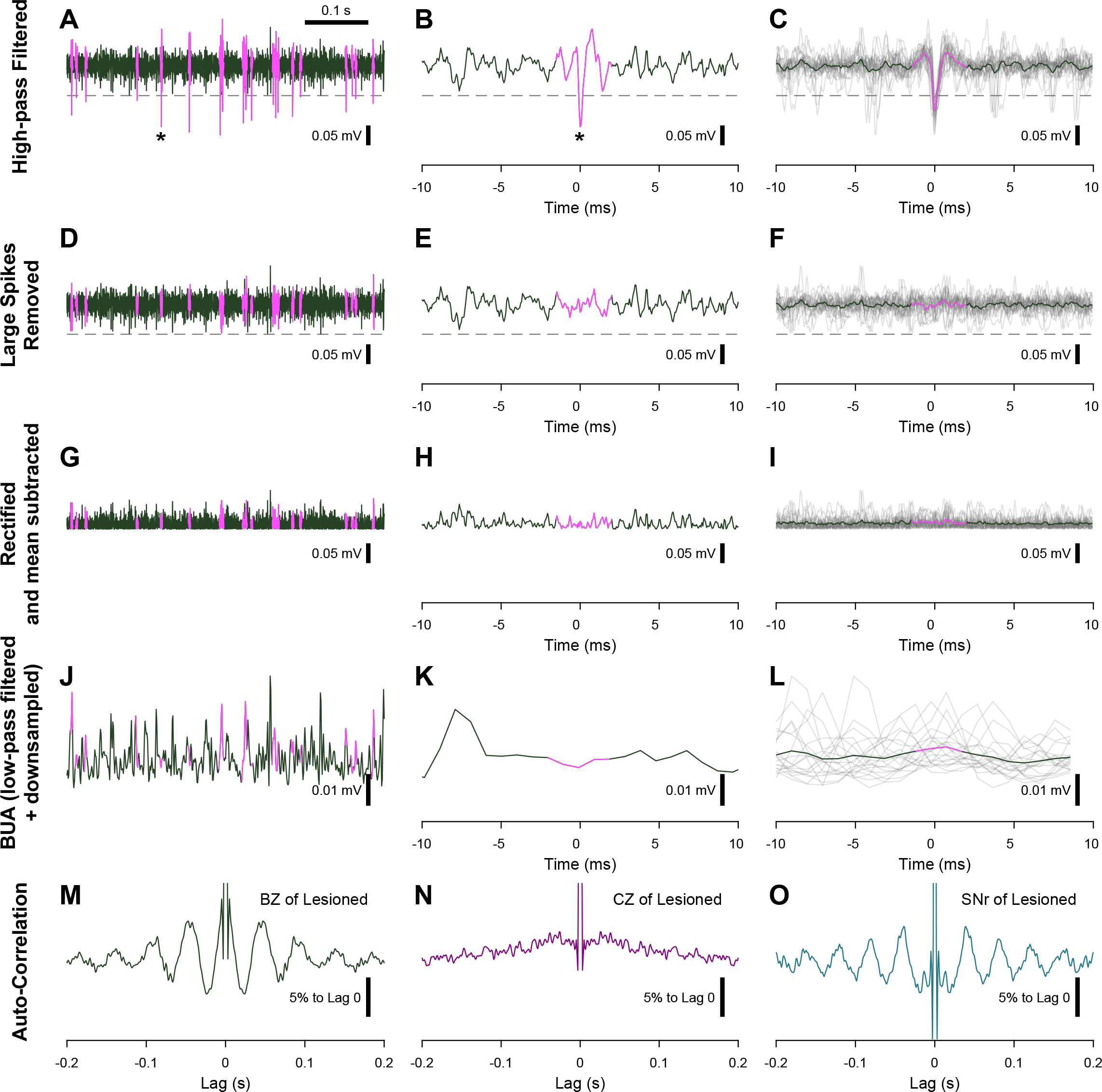
Steps for extraction and processing of background-unit activity (BUA) signals for time series analyses. Wideband recordings made with silicon probes were high-pass filtered at 300 Hz, and then any large-amplitude action potentials (spikes) were detected (***A–C***). Pink spikes in ***A*** are those identified as being of large amplitude (crossing the threshold of 3 standard deviations indicated by the dashed line), with asterisks in ***A*** and ***B*** indicating the same large spike. These large spikes were then removed and replaced with another randomly-selected part of the recording that did not contain large spikes (***D–F***). Replacement data are also highlighted in pink in ***D–L***. The resultant BUA signals were then rectified and mean subtracted (***G–I***) before being low-pass filtered at 300 Hz and downsampled to 1024 Hz to generate a continuous measure for further analyses (***J–L***). Portions of individual signals at each processing step are shown at low (***A***, ***D***, ***G***, ***J***) and high (***B***, ***E***, ***H***, ***K***) temporal resolutions. Several traces are overlaid in relation to the detected large-amplitude spikes to clarify the effects of removing spikes and other signal processing steps (***C***, ***F***, ***I***, ***L***). ***M–O***, Examples of autocorrelation functions of processed BUA signals recorded in the BZ (***M***) and CZ (***N***) of the motor thalamus and in the SNr (***O***) of 6-OHDA-lesioned rats. Note in the BZ and SNr autocorrelations the presence of multiple peaks every 40–50 ms, reflecting oscillations in the beta-frequency band (15–30 Hz).

